# Neuronal exosomes transport a miRNA/RISC cargo to preserve germline stem cell integrity during energy stress

**DOI:** 10.1101/2023.11.15.567290

**Authors:** Christopher Wong, Elena M. Jurczak, Richard Roy

## Abstract

During periods of nutrient scarcity, many animals undergo germline quiescence to preserve reproductive capacity, and neurons are often necessary for this adaptation. We show here that starvation causes the release of neuronal miRNA/Argonaute-loaded exosomes following AMPK-regulated trafficking changes within serotonergic neurons. This neuron-to-germ line communication is independent of classical serotonergic neurotransmission, but instead relies on endosome-derived vesicles that carry a pro-quiescent miRNA cargo to modify germline gene expression. Using a miRNA activity sensor, we show that neuronally-expressed miRNAs can extinguish the expression of germline-mRNA targets in an exosome-dependent manner. Our findings demonstrate how an adaptive neuronal response can change gene expression at a distance by re-directing intracellular trafficking to release neuronal exosomes with specific miRNA cargos capable of tracking to their appropriate destinations.

**One Sentence Summary:** Neurons release miRNA Argonaute-containing exosomes to establish germline stem cell quiescence in response to energy stress.

## Introduction

Despite their shared ability to self-renew following division, many stem cell types remain quiescent unless they receive specific signals to proliferate. This quiescence is likely a conserved means of protecting their invaluable genomic information. We began using *C. elegans* to elucidate the genetic pathways that dictate the various events involved in establishing quiescence in the germline stem cells. During periods of environmental challenge, *C. elegans* larvae can enter a non-ageing “dauer” stage, where development is halted and the germline stem cells arrest (*1*). The AMP-activated protein kinase (AMPK) and its upstream activator, the LKB1 tumor-suppressor orthologue *par-4*, are essential to establish this quiescence. Not surprisingly, mutants that lack all AMPK signalling are completely sterile following recovery from this stage (*2, 3*).

As animals enter the dauer stage, neuronally-expressed AMPK targets a RabGAP protein to ensure that the conserved GTPase RAB-7 remains GTP-bound (*3*). RAB-7 is essential for early-to-late endosome maturation and the generation of exosomes (*4–6*). Recently, this class of extracellular vesicles has attracted attention due to their potential roles in cell-cell communication in both physiological and pathological processes (*7*). However, due to the heterogeneity of extracellular vesicles and the cargos that they transport, it is still unclear whether their release from cells is part of a regulated pathway or indeed a routine shedding of cellular material.

We describe here how the neuronal activation of AMPK in response to starvation results in the formation and secretion of exosomes that carry instructions in the form of miRNAs, from the neurons to the stem cells in the germ line. Following their internalization, these cargo-containing vesicles then transmit a pro-quiescent signal that instructs the germline stem cells to arrest their cell divisions and modify their gene expression to preserve reproductive fitness throughout the course of a potentially lengthy period of energy stress.

### Neuronal RAB-7 localizes to the germ line

We previously identified RAB-7 as a target of TBC-7, a gene that when disabled can suppress the germline defects associated with loss of AMPK signalling (*3*). To determine if RAB-7 is involved in the production of neuronal extracellular vesicles that could transmit information to the germ line during the dauer stage, we expressed GFP::RAB-7 exclusively in the neurons and subsequently imaged dauer larvae. We detected GFP in both the neurons and the germ cells of *daf-2* and *daf-2; aak(0); tbc-7* mutant dauer larvae, but not in *daf-2; aak(0)* (AMPK) mutants (Fig. 1A-C), suggesting that neuronal GFP::RAB-7 localization to the germ line is essential for maintaining post-dauer fertility. The GFP::RAB-7 localization to the perinuclear region is consistent with cells that have accumulated endosomal trafficking components (*8*). No GFP signal could be detected in the germ lines of animals that did not transit through the dauer stage, regardless of their genotype, suggesting that the release of these neuronal GFP::RAB-7-associated structures is specific to this physiological response (fig. S1A-C). The GFP::RAB-7 signal was still visible in the germ cells of *daf-2; aak(0); tbc-7* mutant animals that recovered from this stage (fig. S1D, F), while no GFP signal was detected in post-dauer *daf-2; aak(0)* mutants (fig. S1E). To examine if this mechanism of neuron to germ line communication is conserved in other modes of dauer entry, neuronal GFP::RAB-7 was expressed in *daf-7* and wild-type N2 animals treated with dauer pheromone and image analysis was performed on the resultant dauer larvae. GFP signal was detected in the germ cells of both *daf-7* and N2 dauer larvae, suggesting that the transfer of neuronal GFP::RAB-7 to the germ cells at the onset of dauer development is a general, conserved feature that is shared among the best characterized pathways responsible for dauer formation (fig. S2A-B).

**Fig. 1.**
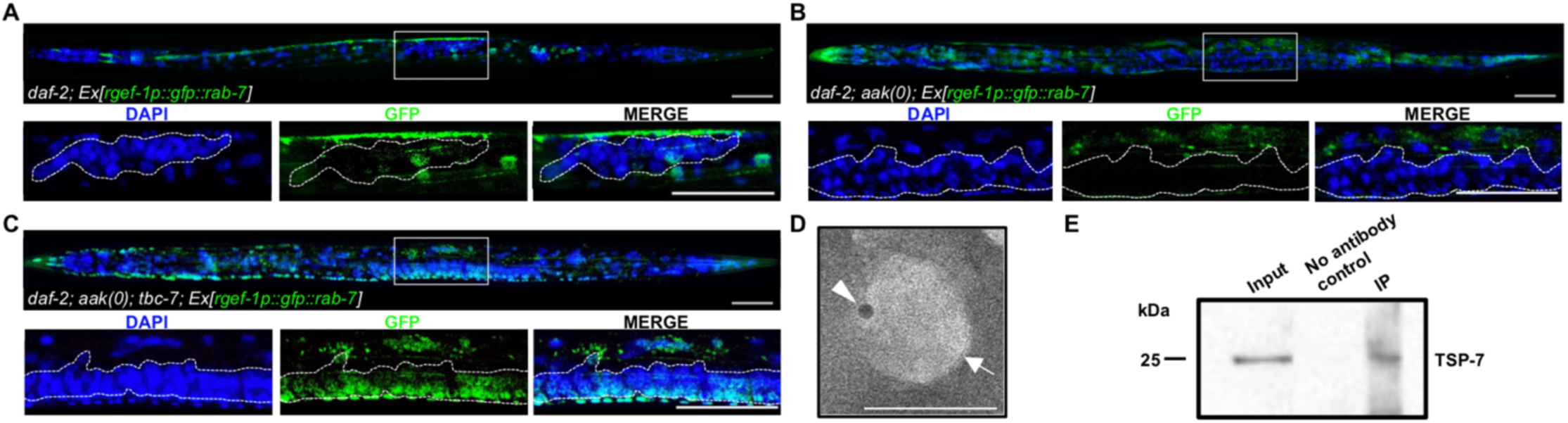
Neuronal RAB-7 is associated with exosomes that localize to the germ line during energy stress. (**A-C**), Confocal images of neuronal GFP::RAB-7 expression in dauer larvae in (A) *daf-2*, (B) *daf-2; aak(0)*, and (C) *daf-2; aak(0); tbc-7* mutants. Solid white frame indicates position of the higher magnification insets (below). Dotted white lines outline the gonad. Scale bars, 50 !m. Representative of three independently-generated transgenic lines. (**D**) Negative stain TEM on isolated exosomes from *daf-2* mutants that express neuronal HA::RAB-7 with 6 nm immunogold against HA. White arrowhead indicates gold particle bound to exosome. White arrow indicates lipid bilayer. Scale bar, 50 nm. (**E**) Immunoprecipitation was performed on pan-exosomes isolated from *daf-2* mutants expressing neuronal HA::GFP::RAB-7. Western analysis with anti-GFP antibodies was conducted against pan-exosome input, no antibody control, and the immunoprecipitation against HA::GFP::RAB-7 using antibodies against HA. Representative of two independent experiments.

To determine if the localization of GFP::RAB-7 to the germ line is a result of classical neurotransmitter release, we assessed the post-dauer fertility of *daf-2; aak(0); tbc-7* mutants with either compromised synaptic vesicle or dense-core vesicle secretion (*9, 10*). The post-dauer fertility was unaffected by these mutations (fig. S3A), suggesting that these GFP::RAB-7 structures do not stem from vesicle-mediated peptide secretion or synaptic vesicle release. Due to the previously described role of RAB-7 in endosome to lysosome trafficking and autophagy (*11*), we also determined whether autophagy was affected in mutants with altered RAB-7 activity (*daf-2; aak(0); tbc-7*). As autophagy progresses, the *C. elegans* orthologue of Atg8/LC3, LGG-1 is lipidated to become LGG-1 phosphatidylethanolamine (PE), then subsequently cleaved to eventually contribute to the formation of the autolysosome (*11*). Using a GFP-tagged LGG-1 transgene, we assessed autophagic flux through Western analysis to conclude that autophagy is unaffected in *tbc-7*-compromised mutants (fig. S4A).

Because RAB-7 functions both in endosomal maturation and exosome production, we characterized these GFP-containing structures by immunogold labelling TEM following their purification from dauer larvae. GFP::RAB-7 localized to the membrane of these isolated vesicles, which were 30-150 nm, within the diameter range typical of small extracellular vesicles (sEVs) derived from the endosomal pathway (*12*) (Fig. 1D). To assess if these sEVs could be exosomes, we performed immunoprecipitation using an anti-HA antibody against HA::GFP::RAB-7 on the vesicle sample and performed Western analysis against TSP-7, the *C. elegans* orthologue of CD63, a marker for exosomes derived from the endosomal route (*13*). Our Western analysis indicated that TSP-7 was indeed associated with our GFP-containing RAB-7 structures consistent with the possibility that these vesicle-like structures could indeed be exosomes derived from the endosomal pathway (Fig. 1E, fig. S5A). Our data therefore support a role for RAB-7 in contributing to the formation of specialized neuronal vesicles following AMPK activation (*3*). These RAB-7-associated exosomes exit the neurons and thereafter translocate to the germ cells during the dauer stage, where RAB-7 remains up to 24h after recovery from the dauer stage.

In *C. elegans*, the nervous system uses an array of classical neurotransmitters similar to mammalian models in order to influence the activity of molecular pathways in post-synaptic neurons. Since these neurons have specialized functions in signalling, it is possible that *tbc-7* functions in one class of neurons that is critical for responding to the energy stress that is associated with dauer formation. To determine which neurons *tbc-7* functions in, we drove the expression of wild-type *tbc-7* under neuron type-specific promoters in *daf-2; aak(0); tbc-7* mutants and assessed the resultant post-dauer fertility of the transgenic mutants, whereby the re-introduction of a wild-type copy of *tbc-7* in the relevant neurons should restore the wild-type function of the RabGAP and render the post-dauer animals sterile. The expression of wild-type *tbc-7* exclusively in serotonergic neurons (*tph-1p*) reverted the suppression of AMPK germline defects (fig. S6A) and driving the expression of GFP::RAB-7 in the serotonergic neurons was sufficient to detect GFP signal in the germ cells of the transgenic dauer larvae (fig. S6B). These data suggest that the serotonergic neurons could be the major, but potentially not the only, class of neurons that produce and secrete exosomes in response to energy stress through the activity of the AMPK/TBC-7/RAB-7 signalling axis during the dauer stage.

Because we observed neuronally-derived GFP::RAB-7 in the germ cells we questioned whether RAB-7 alone was sufficient to navigate all RAB-7-associated exosomes to the germ cells. We therefore expressed GFP::RAB-7 in multiple cell types using tissue-specific promoters but despite visible GFP signal in each of the tissues assayed, no GFP could be detected in the germ cells of dauer larvae (fig. S7A-C). These data indicate that although neuronal RAB-7 is critical for the formation and release of these exosomes, it does not function alone. An additional navigator component must be required to direct the neuronal RAB-7-associated exosomes to the germ line.

### miRNAs regulate germline gene expression

The neuronal RAB-7-associated exosomes could provide a pro-quiescent signal that instructs the stem cells to modify their chromatin and adjust their gene expression in anticipation of a potentially lengthy environmental challenge. We have shown that small RNA pathways are important effectors downstream of AMPK (*2, 14–16*), while miRNAs and other RNAs have been identified in the cargo of several types of exosomes (*17*). Therefore, to determine if the AMPK-dependent germline quiescence is regulated through a small RNA-based mechanism, we soaked *daf-2; aak(0)* mutants in a solution that contained RNA extracted from *daf-2* dauer larvae (see Materials and Methods). Surprisingly, this RNA was sufficient to suppress the post-dauer sterility of the AMPK mutants (Fig. 2A), indicating that one or more RNA species in this mixture acts downstream of AMPK and was capable of restoring quiescence to the germline stem cells in the AMPK animals. Furthermore, this RNA-mediated suppression did not require the miRNAs *mir-1* and *mir-44*, both of which have been shown to regulate *tbc-7* activity during the dauer stage (*3*) (Fig. 2A).

**Fig. 2.**
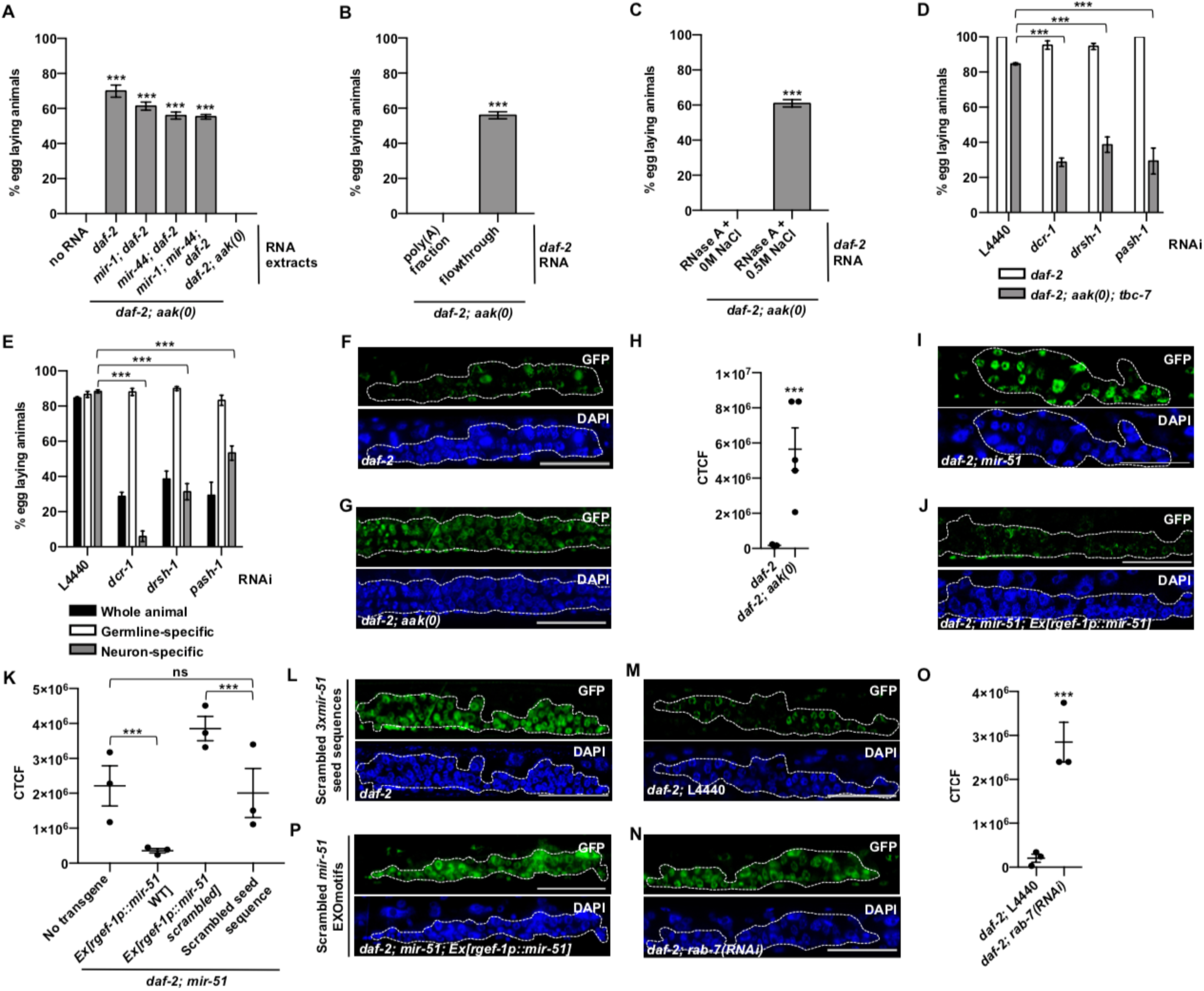
Neuronally-expressed miRNAs act non-autonomously to preserve germ cell integrity. (**A-C**) Small dsRNAs affect post-dauer fertility of *daf-2; aak(0)* mutants soaked in (A) RNA extracted from animals of indicated genotypes, (B) RNA extracted from *daf-2* dauer larvae that were treated with RNase A in either 0 M or 0.5 M NaCl, and (C) either the poly(A) fraction or a poly(A)-depleted fraction (flowthrough) of total RNA extracted from *daf-2* mutants. (**D-E**) Post-dauer fertility of *daf-2; aak(0); tbc-7* mutants fed with bacteria expressing dsRNA that corresponds to components involved in miRNA biogenesis in animals subjected to (D) whole animal RNAi or (E) tissue-specific RNAi. (**F-G**) Confocal images of the miRNA sensor containing three wild-type *mir-51* seed sequences in (F) *daf-2* and (G) *daf-2; aak(0)* dauer larvae. (**H**) Corrected total cellular fluorescence (CTCF) of GFP intensity by genotype. (**I-J**) Confocal images of the miRNA sensor containing three wild-type *mir-51* seed sequences in (I) *daf-2; mir-51* and (J) *daf-2; mir-51* that express a *mir-51* transgene driven by a pan-neuronal promoter (*rgef-1p::mir-51*) dauer larvae. (**K**) Corrected total cellular fluorescence (CTCF) of GFP intensity by genotype. (**L**) Confocal image of the miRNA sensor containing three scrambled *mir-51* seed sequences in *daf-2* dauer larvae. (**M-N**) Confocal images of the *mir-51*-specific miRNA sensor in *daf-2* dauer larvae fed with bacteria that express (M) empty vector L4440 or (N) dsRNA that corresponds to *rab-7*. (**O**) Corrected total cell fluorescence (CTCF) of GFP intensity by genotype and RNAi condition. (**P**) Confocal image of the miRNA sensor in *daf-2; mir-51* mutants expressing neuronal *mir-51* with three scrambled EXOmotifs. Dotted white lines outline the gonad. Scale bars, 50 !m. Data is mean ± SEM. *P* values are derived from Marascuilo procedure (A-E) and one-way ANOVA (H, K, O). Post-dauer fertility data are representative of three independent experiments; n = 50 animals per condition for post-dauer fertility. Confocal imaging and CTCF data is representative of three independent experiments; n = 15 animals per condition. \*\*\**P* < 0.0001. ns, not significant.

To examine if the suppression of the germline defects might act through an mRNA-based mechanism, we repeated the RNA soaking experiment, but prior to soaking, we separated the RNA mixture using oligo d(T)_25_ beads to create one fraction with only mRNA and another that was poly(A) mRNA depleted (flowthrough). When the *daf-2; aak(0)* animals were incubated with the mRNA fraction, these animals remained post-dauer sterile (Fig. 2B). However, when the *daf-2; aak(0)* mutants were incubated with the flowthrough fraction containing all small non-coding RNAs (Fig. 2B), the animals were post-dauer fertile.

To test further what RNAs might be involved in the RNA-mediated restoration of germline competence, we digested the RNA extracts with RNase A. RNase A digests all RNA efficiently at 0 M NaCl, but at 0.3 M NaCl or higher, RNase A selectively digests single-stranded RNA, leaving double-stranded RNAs intact (*18, 19*). When *daf-2; aak(0)* mutants were incubated with RNA previously treated with RNase A without NaCl, the animals were completely sterile (Fig. 2C). However, when the RNA solution was treated with RNase A in 0.3 M NaCl prior to incubation with *daf-2; aak(0)* mutants, the post-dauer sterility was completely suppressed (Fig. 2C). These data suggest that the rescuing component of our RNA solution was not mRNA, but was more likely a double-stranded RNA(s) that can be processed into small RNAs. These would then be responsible for the maintenance of germ cell quiescence/integrity during the dauer stage.

From these and other data, we know that small RNAs affect germline quiescence (*2, 3*), but whether the role of small RNAs could be dependent on the release of neuronal exosomes has not been established. By performing directed or unbiased RNAi surveys in fertile AMPK mutant animals that have activated RAB-7 (*daf-2; aak(0); tbc-7)*, we sought to identify components that are required for this process; gene products potentially involved in a) the production and trafficking of the exosomal cargo, b) transporting the exosomes between tissues, and even c) effecting the chromatin changes in the germline stem cells.

DCR-1 is an endoribonuclease III-like enzyme that is essential for the production of all small RNAs, with the exception of piRNAs (*20*). RNAi of *dcr-1* in *daf-2; aak(0); tbc-7* mutants resulted in post-dauer sterile animals (Fig. 2D), indicating that it, and its small RNA products, are required for germline integrity during this stage. We then performed RNAi to compromise each of the hingepin biosynthetic enzymes for every class of small RNAs that require *dcr-1* activity. From this analysis we noted that when the components required for miRNA biogenesis were disabled, the *daf-2; aak(0); tbc-7* mutants became post-dauer sterile, while the loss of other factors required for siRNA production or function had no effect on post-dauer fertility (Fig. 2D, fig. S8A), suggesting that only miRNAs are involved in preserving germ cell integrity during or after the dauer stage.

Because AMPK and TBC-7/RAB-7 function together in the neurons to regulate germline integrity during the dauer stage, we used a previously verified tissue-specific RNAi strategy to confirm where miRNA biogenesis is required to provide the pro-quiescent signal to the germline stem cells (*3*). Our findings indicated that when miRNA biogenesis was compromised exclusively in the neurons, we reproduced the same post-dauer sterility described above, while disabling these same factors in the germ line had no effect (Fig. 2E). These data are consistent with a model where miRNAs that are produced in the neurons are responsible for the gene expression changes required in the germline stem cells downstream of RAB-7 activation.

To evaluate if the neuronally-produced miRNAs could affect gene expression in the germ line through a RAB-7-dependent mechanism, we developed a miRNA activity sensor that would be responsive to neuronally-expressed miRNAs. We recently revealed that the levels of many miRNAs are affected by the loss of AMPK (*21*). Among them, *mir-51* seems to be among the most significantly affected (*21*). However, when *mir-51* levels were increased in the neurons, post-dauer fertility was restored significantly (fig. S9A). Using a fluorescent germline miRNA activity sensor created by driving HIS-15::GFP that contains three binding sites for *mir-51* in its 3’UTR (hereafter referred to as “sensor”), we tested whether the GFP (sensor) could be silenced in a *mir-51*-dependent manner (fig. S9B). In replete conditions, GFP was expressed at similar levels in the adult stage in both *daf-2* and *daf-2; aak(0)* mutants (fig. S9C-E). However, during the dauer stage, *daf-2* animals silenced the sensor in the germ cells, while the expression of the sensor was unaffected in dauer larvae that lacked AMPK (Fig. 2F-H). To show that this effect is specific to the activity of *mir-51*, we expressed *mir-51* exclusively in the neurons of *mir-51* deletion mutants and found that the transgenic mutant animals nevertheless silenced the sensor (Fig. 2I-K). Next, we scrambled the *mir-51* seed sequences in the sensor of *daf-2* animals, preventing *mir-51* from binding to the 3’UTR (fig. S9B) (see Methods and Materials). These sequence changes in the *mir-51* target site resulted in the full restoration of HIS-15::GFP expression in the germ cells (Fig. 2L, K). Taken together, these data suggest that miRNAs, such as *mir-51*, that are expressed in the neurons, can silence germline mRNA targets that possess its target site, and that this regulation is both specific and AMPK-dependent during the dauer stage.

Because neuronally-expressed *mir-51* can act cell non-autonomously to silence germline targets and RAB-7-containing exosomes exit the neurons to ensure germline quiescence, it seemed plausible that these exosomes carry a miRNA cargo that provides a pro-quiescent signal to the germline stem cells in anticipation of the dauer stage. To determine if the silencing of the sensor we observed with *mir-51* involves the RAB-7-associated exosomes, we compromised the production of exosomes by RNAi and compared the levels of the sensor with that of control animals. Inhibiting the production of exosomes by *rab-7* RNAi completely abrogated the silencing of the sensor by *mir-51*, while empty vector RNAi controls remained fully capable of silencing the GFP expression (Fig. 2M-O). Therefore, the neuronal miRNA-dependent silencing of germline targets we observed in dauer larvae is entirely dependent on *rab-7*, most likely through its role in the formation of exosomes.

The loading of miRNA cargos into exosomes relies on a number of factors, including short nucleotide stretches called EXOmotifs that act as sorting sequences for miRNAs and determine whether the miRNA will be retained within the cell or packaged into exosomes (*22*). Any disruption of these motifs results in the exclusion of the miRNA from exosomes (*22*). *mir-51* possesses three of these motifs, consistent with its potential incorporation in RAB-7-associated exosomes. To examine if these EXOmotifs contribute to the sorting of neuronally-produced *mir-51* into exosomes to target the germ line, we expressed a *mir-51* variant with scrambled EXOmotifs in the neurons of animals that lack *mir-51* and assessed its ability to silence the sensor. *In silico* modeling predicts that scrambling of the EXOmotifs in *mir-51* would not affect its synthesis or secondary structure (fig. S9F, G), yet the scrambled mutant *mir-51* variant failed to silence the germline sensor (Fig. 2K, P). Consistent with our imaging results, mutants with the scrambled EXOmotif *mir-51* variant could no longer suppress the post-dauer sterility, suggesting that *mir-51* is likely to be sorted into exosomes through the recognition of its EXOmotifs in order to ultimately reach its germ line targets.

In addition to showing that a neuronally produced miRNA can regulate an artificially expressed germline sensor, we wanted to demonstrate that it could regulate the germline chromatin landscape and subsequent gene expression. To show that these neuronally-derived exosomes play a role in regulating the germline gene expression program during the dauer stage, we first assessed the levels of chromatin marks in *daf-2; aak(0)* mutants expressing neuronal *mir-51*. In *daf-2; aak(0)* dauer larvae with no additional transgenes, the levels of all the chromatin marks we assessed were upregulated (*2*). However, in *daf-2; aak(0)* mutants expressing neuronal *mir-51*, the abundance of both activating and repressive chromatin marks were partially restored to near wild-type levels (fig. S10A-B). Next, to examine if the abnormal gene expression caused by the aberrant chromatin modifications in *daf-2; aak(0)* mutants were also corrected, we performed RT-qPCR to assess the gene expression of *daf-2; aak(0)* mutants expressing neuronal *mir-51*. We selected germline-expressed genes that are differentially expressed during the dauer stage in *daf-2; aak(0)* mutants as compared to *daf-2* mutants (*2, 23*). RT-qPCR revealed that the expression of these selected germline genes in *daf-2; aak(0)* mutants expressing neuronal *mir-51* largely resemble the expression patterns of *daf-2* mutants (fig. S10C), suggesting that the neuronal expression of *mir-51* affects germline gene expression during the dauer stage. Altogether, our data suggest that miRNAs synthesized in the neurons are incorporated into RAB-7 exosomes, likely through the presence of *cis* acting EXOmotifs. They are then released, and subsequently make their way to the germ line where they adjust gene expression to promote quiescence in the germline stem cells.

### Small RNA regulators act at distinct stages

To identify additional components involved in the transport of the exosomal miRNAs from the neurons to the germ line, we employed a hypomorphic RNAi strategy in *daf-2; aak(0); tbc-7* (AMPK mutant animals with active RAB-7) to survey the effects of compromising individual genes involved in dsRNA transport on post-dauer fertility. RNAi of *sid-3*, *sid-5*, and *rme-8* in these mutants reduced post-dauer fertility significantly, indicating that their effects are required for the appropriate changes in germline gene expression mediated by the RAB-7-associated exosomes (Fig. 3A). SID-5 is a single-pass transmembrane RNA channel and has been shown to localize with endosomal components like RAB-7 and LMP-1, thus linking RNA mobility to vesicular transport (*24*). RME-8 is a conserved protein required for endocytosis, and similarly to SID-5, localizes to endosomal membranes (*25*). Because SID-5 has been shown to function in a cell non-autonomous manner as a dsRNA exporter rather than importer (*24*), we examined in which tissues these RNA transport proteins could be required by using strains where RNAi activity is restricted to either the neurons or the germ line (*24, 25*). These RNAi experiments indicated that *sid-5* and *rme-8* are required exclusively in the neurons, consistent with their known roles in RNA export and endocytic maturation, respectively (Fig. 3B). On the other hand, the conserved non-receptor tyrosine kinase *sid-3,* was required in the germ line (Fig. 3B). SID-3 is required for the efficient uptake of dsRNA in *C. elegans* (*26*), while its mammalian ortholog (ACK) was initially identified as a binding partner of Cdc42, a small GTPase that promotes endocytosis (*27*). A kinase-dead SID-3 variant is unable to import dsRNA into *C. elegans* cells (*26*). Therefore, to examine if the kinase domain of SID-3 is required for the maintenance of germ cell integrity, we expressed a kinase-dead variant of SID-3 in the *daf-2; aak(0); tbc-7* mutants (*26*). Unlike animals with wild-type SID-3, the *daf-2; aak(0); tbc-7* mutants that expressed a kinase-dead SID-3 variant could no longer suppress the germline defects associated with a lack of AMPK signaling during the dauer stage or following recovery (Fig. 3C, D). Moreover, mutants with compromised *sid-3* expression or a kinase-dead SID-3 variant blocked the miRNA-dependent silencing of the miRNA sensor in the germ cells (fig. S9I-K), similar to *daf-2; aak(0)* mutants.

**Fig. 3.**
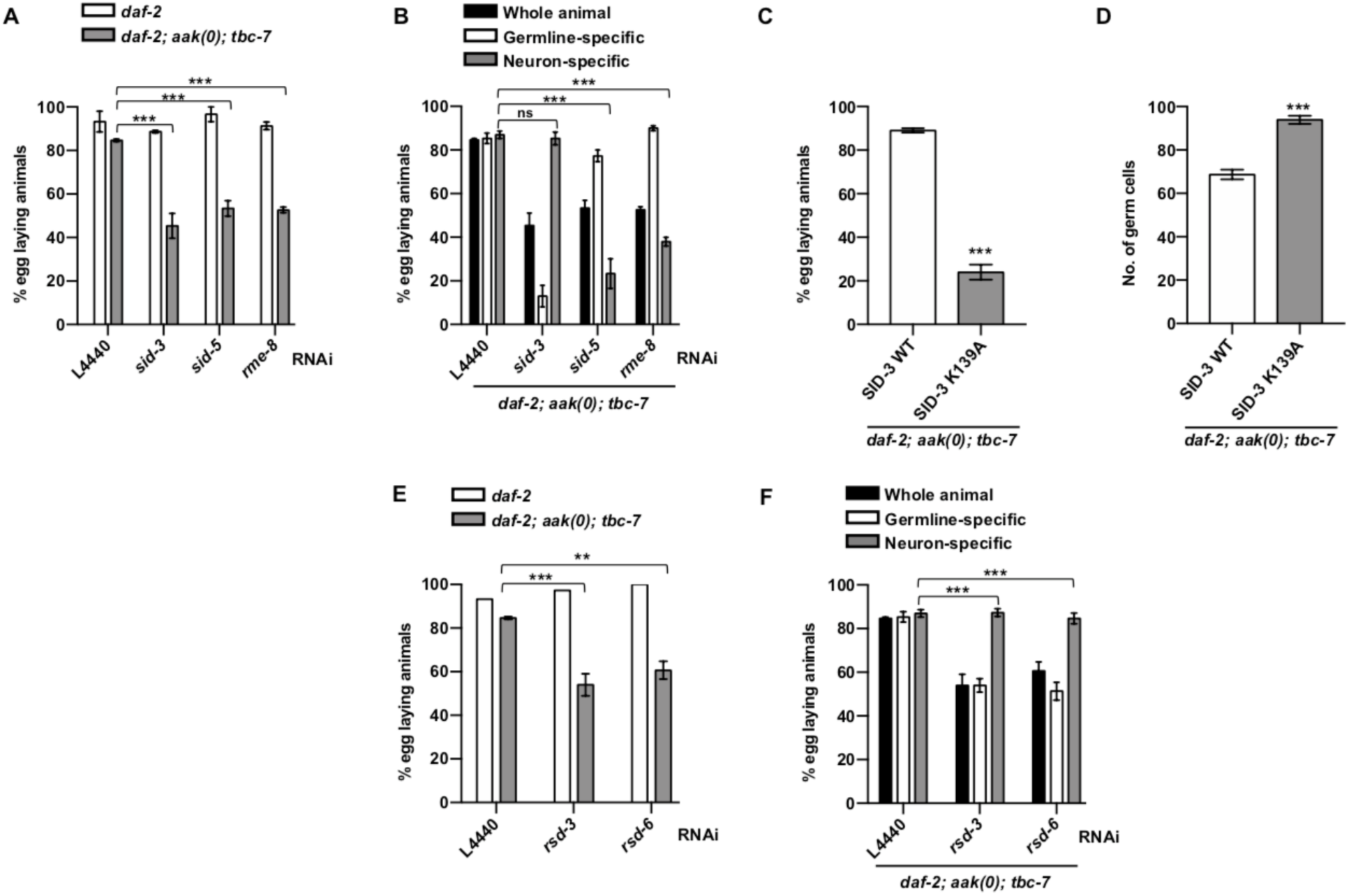
miRNA-containing exosomes require dsRNA transport proteins to efficiently transmit pro-quiescent signals to the germline stem cells. (**A-B**) Post-dauer fertility of *daf-2; aak(0); tbc-7* mutants fed with bacteria expressing dsRNA that corresponds to various small RNA transporters in animals subjected to (A) whole animal RNAi or (B) tissue-specific RNAi. (**C-D**) The (C) post-dauer fertility and (D) number of germ cells in the dauer larvae of *daf-2; aak(0); tbc-7* mutants with a kinase-dead variant of SID-3. (**E-F**) Post-dauer fertility of *daf-2; aak(0); tbc-7* mutants fed with bacteria expressing dsRNA that corresponds to RNAi spreading components in animals subjected to (E) whole animal RNAi or (F) tissue-specific RNAi. Data is mean ± SEM. *P* values are derived from Marascuilo procedure (A, B, C, E, F) or ordinary one-way ANOVA (D); n = 50 animals per condition for post-dauer fertility. n = 25 animals per condition for number of germ cells. Post-dauer fertility and number of germ cells data are representative of three independent experiments. \*\*\**P* < 0.0001.

Following the incorporation of the exosomes, the germ cells undergo a concerted cell cycle arrest associated with a general modification of the chromatin (*2*). This invokes the possibility of an amplification mechanism, whereby the pro-quiescent instructions conveyed by the miRNAs must be uniformly distributed throughout the entire germ line. The RNAi spreading defective (*rsd*) genes have been shown to regulate both the import of dsRNA into cells and the propagation of the small RNA-based signals (*28, 29*). *rsd-2* and *rsd-3* encode proteins that associate with intracellular trafficking components and endomembranes, where the loss of either one of these gene products results in silencing defects, not only in the tissue where the dsRNA is acting, but also in adjacent cells and tissues (*29*). Consistent with their implication in this endosome-mediated process, the loss of *rsd-3* and *rsd-6* expression resulted in extensive post-dauer sterility, while further interrogation indicated that these two genes are only required in the germ line (Fig. 3E, F). Like SID-3, RSD-3 has a conserved ENTH domain that is required for RNA import into somatic and germ cells (*29*), and although RSD-6 does not have an ENTH domain, it does possess a Tudor domain, which has known roles in transposon silencing through RNAi activity (*28, 30, 31*). Therefore, like SID-3, RSD-3 and RSD-6 could be required for exosomal miRNA uptake and/or to propagate or amplify small RNA signals uniformly throughout the germ line.

### Neuronal miRISC translocates to the germ line with RAB-7

The neuronal exosomes could potentially transport pre-miRNAs, which would require assembly into a miRISC complex in the germ cells, perhaps requiring the function of the *rsd* genes. Alternatively, the miRNAs could be expressed and pre-assembled with their cognate miRISC Argonaute proteins poised to act upon delivery. To distinguish between these possibilities, we expressed the miRNA Argonautes ALG-1 and ALG-2 fused to mKate in the neurons and imaged the transgenic dauer larvae to assess whether the Argonautes remained within the neurons, or like the RAB-7-containing exosomes, they could be detected in both the neurons and in the germ cells. For both ALG-1 and ALG-2, fluorescent signal was detected in the germ cells of *daf-2* and *daf-2; aak(0); tbc-7* mutants (fig. S11A, C, E), but not in the germ lines of *daf-2; aak(0)* mutants or mutants expressing neuronal mKate alone, thus exhibiting a similar expression pattern to GFP::RAB-7 (Fig. 4A, fig. S11B, D, F). Furthermore, the localization of ALG-1/-2 mKate signal in the germ line was dependent on *rab-7*, although the mKate signal remained strong in the neurons of the *rab-7(RNAi)* animals (Fig. 4B, C).

**Fig. 4.**
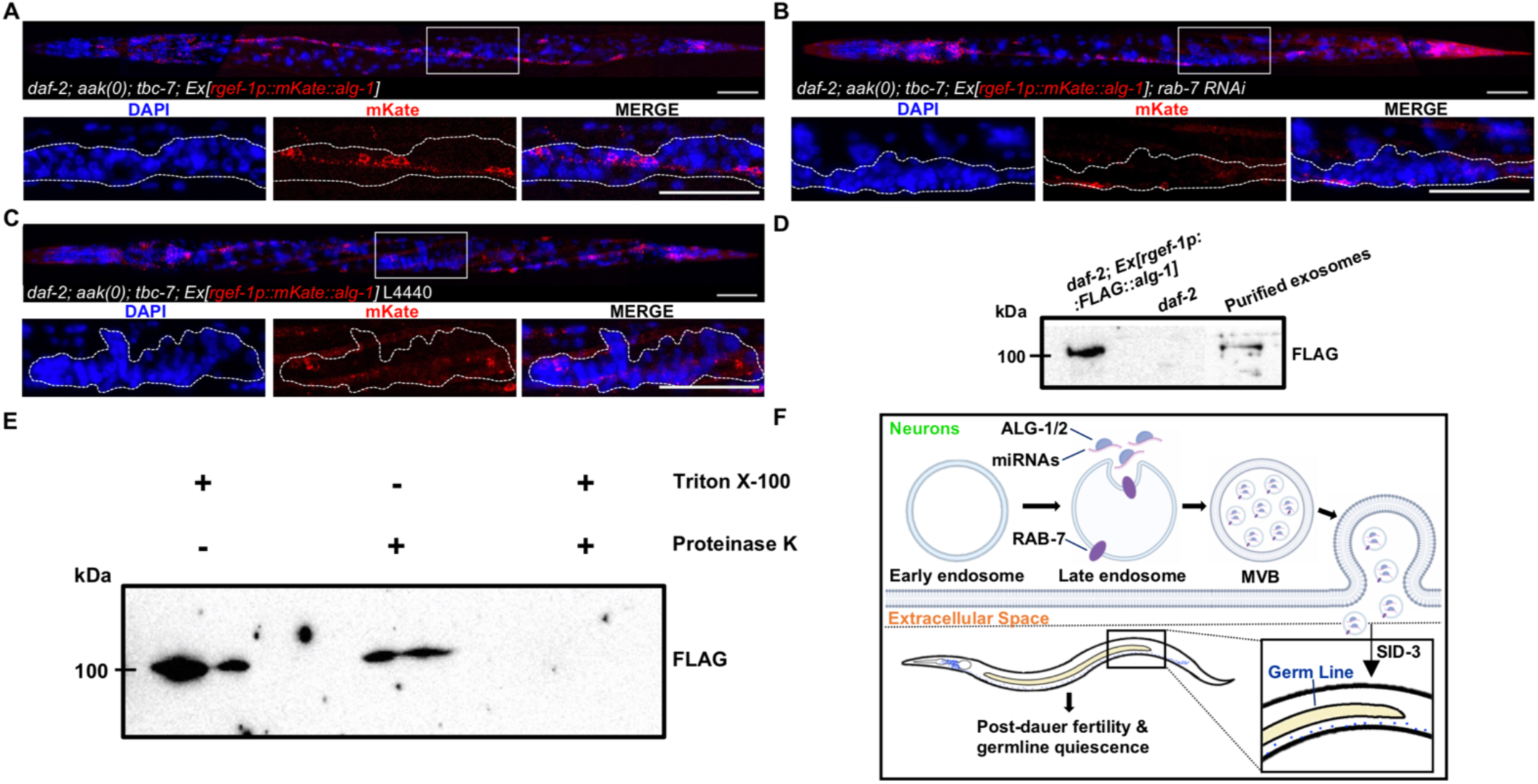
miRNA Argonautes are incorporated into neuron-derived exosomes and subsequently localize to the germ line. (**A**) Confocal images of neuronal mKate::ALG-1 expression in *daf-2; aak(0); tbc-7* dauer larvae. (**B-C**) Confocal images of neuronal mKate::ALG-1 expression in *daf-2; aak(0); tbc-7* dauer larvae fed with bacteria that express (B) empty vector L4440 or (C) dsRNA that corresponds to *rab-7*. Solid white frame indicates position of the higher magnification insets (below). Dotted white lines outline the gonad. Scale bars, 50 !m. Representative of three independently generated transgenic lines. (**D**) Western analysis performed with anti-FLAG antibodies on exosomes isolated from mutants expressing neuronal FLAG::ALG-1. Western blots were spliced together due to differences in exposure times. (**E**) Western analysis performed with anti-FLAG antibodies on exosomes isolated from mutants expressing neuronal FLAG::ALG-1 that have been treated with either Triton X-100, Proteinase K, or both. Representative of two independent experiments. (**F**) Proposed model of the secretion of neuronal exosomes containing miRNAs to protect germline integrity.

If the Argonautes are complexed with their miRNA cargo within exosomes they should also purify with our exosome preparations. Western analysis indicated that ALG-1 was present in samples of purified exosomes (Fig. 4D). Moreover, ALG-1 is associated with its miRNA cargo on the interior of the exosome and not non-specifically bound to the exosomal outer surface. When the exosomes were treated with Proteinase K alone, we were still able to detect ALG-1, indicating that the lipid bilayer of the exosome protects ALG-1 from proteolytic degradation. However, when the exosomes were treated with detergent, then with the protease, ALG-1 was no longer detectable (Fig. 4E). This indicates that the solubilization of the exosomal membrane permitted Proteinase K to digest ALG-1 that was initially protected by the lipid barrier. These data suggest that ALG-1 is incorporated into the neuronal exosomes, potentially bound to its miRNA cargo, in a regulated manner. The miRISC complex is then released from the neurons in the RAB-7-containing vesicles; these entities home specifically to the gonad, where following internalization, the miRNAs can instruct the germline stem cells to execute quiescence in response to the environmental cues associated with dauer development.

## Discussion

The emergence of exosomes as critical extracellular agents capable of delivering biologically relevant signalling molecules to target tissues has been fueled by numerous studies in various models. Their purification and subsequent analysis have indicated that these vesicles carry a diverse macromolecular cargo that can include DNA, proteins, mRNA, miRNAs, and other RNA fragments (*17, 22, 32*). Yet, despite the identification of these important molecules, the effects of releasing intracellular contents to neighbouring cells or tissues have yet to be fully elucidated due to the large heterogenicity of extracellular vesicles and their many biogenesis and loading pathways. While neurons and neuronal small RNAs have been shown to regulate changes in germline gene expression in *C. elegans*, a genetic, mechanistically defined pathway that could account for this intertissue communication has not been described.

Using an unbiased genetic approach, we have shown how the formation and release of RAB-7-containing exosomes from the neurons is required to signal non-autonomously to the *C. elegans* germ line (Fig. 4F). These exosomes carry pro-quiescent instructions in the form of miRNAs that are required to adjust gene expression in the germline stem cells to block their cell divisions, while preparing the genome for an extended period of stress that would otherwise compromise reproductive fitness.

The extensive energy stress associated with starvation could presumably affect the levels of critical macromolecular building blocks, and/or the energy-dependent quality control mechanisms that monitor their synthesis in the germ cells. That the serotonergic neurons have evolved a role in adjusting germline gene expression in response to energy stress via AMPK activation is a particularly efficient means of protecting genome integrity in the “immortal” germ lineage. These neurons are highly sensitive to intracellular changes in energy levels, eliciting adaptive modifications to foraging behaviours to correct any perceived energy deficit (*33*).

Although the function of AMPK in this subclass of neurons has already been described for its role in behavioral modification in response to starvation (*34, 35*), our data indicate that AMPK activation in the serotonergic neurons acts at multiple levels. In addition to its critical role in altering feeding strategies, its activation will also initiate a specific trafficking process that culminates with the incorporation of specific miRNAs, among probably additional molecules, into RAB-7-associated exosomes earmarked for release and targeting to the germ line.

Using our miRNA sensor, we confirmed the importance of EXOmotifs in the packaging of miRNAs into exosomes, while also demonstrating that their expression in neurons results in the efficient silencing of their cognate transcripts in a distant tissue. Whether this requires nascent transcription of EXOmotif-harbouring miRNAs remains to be resolved, but either the expression or the selection of these transcripts is dependent on neuronal AMPK. Alternatively, perhaps all EXOmotif-containing miRNAs and their bound Argonautes are indiscriminately incorporated into the maturing neuronal exosomes. The exosomes would then carry a constant miRNA cargo, and following internalization, only relevant mRNA targets expressed in the target tissue would be affected. In this scenario, the AMPK-dependent activation of RAB-7 would be sufficient to initiate the process.

The molecular basis of the germline translocation of the exosomes and the role of the dsRNA transporter SID-3 in their internalization is still unclear. Nevertheless, our genetic analysis indicates that the miRNA-containing exosomes access the germline stem cells to effect changes in their gene expression. However, to ensure that all the germline stem cells respond uniformly, the miRNA signal must be amplified and spread throughout the entire germ line. This is not consistent with the current model of miRNA action, whereby miRNAs act stoichiometrically. It therefore invokes the possibility of an intermediate that is responsible for either amplifying or relaying the signal throughout the germ line.

The pro-quiescent function of these exosomes relies on RAB-7 to generate the appropriate vesicles, but other factors provide the navigational function responsible for germline homing. In the developmental context we describe here, the exosomes deliver pro-quiescent instructions in the form of miRNAs to the germline stem cells, but it is unlikely that such an elaborate mechanism of cell non-autonomous regulation would be exclusive to the dauer stage. It is most probable that this mechanism is used in other physiological situations in *C. elegans* and/or in other organisms. By exploiting the EXOmotif-based incorporation into these vesicles, virtually any miRNA, or potentially any RNA, may be incorporated into these vectors for precise tissue-specific targeting. The key factors involved in directing these vesicles to desired tissue locations will ultimately come from purification and characterization of the neuronal exosome-associated constituents that are responsible for the navigator function.

Our work highlights the function of these neuronal exosomes in the epigenetic modification of germ cell gene expression. Although we did not test if or how these vesicles might contribute to any transgenerational epigenetic phenomena, past work has shown that neuron-specific small RNAs communicate with the germ line to adaptively modify behaviours transgenerationally (*36*). Curiously, in the late 19th century Darwin proposed a role for “gemmules” in the transmission of heritable traits (*37*). Our findings underscore a role for neuronal vesicles that bring about epigenetic change in the germ line to adapt gene expression to protect the reproductive competence of the germ cells. We do not yet know if these, or related vesicles, might also act as vectors to convey heritable epigenetic modifications, but if that bears true, Darwin’s “gemmule” hypothesis may not have been so far removed from the actual mechanisms that drive these changes.

## Acknowledgments

The Canadian Institutes of Health Research (Project Grant, CIHR PJT-180267) supported this work. We thank the *Caenorhabditis elegans* Genetics Center (CGC) for *C. elegans* strains. We thank the Facility for Electron Microscopy Research at McGill University for help with sample preparation and microscope operation. We thank Monique Zetka for providing the anti-HA antibody. We thank Shaolin Li and Ryan Dawson for technical assistance.

## Funding

The Canadian Institutes of Health Research Project Grant CIHR PJT-180267 (RR)

## Author contributions

Conceptualization: CW, RR Methodology: CW, RR Investigation: CW, EMJ Visualization: CW

Funding acquisition: RR Supervision: RR

Writing – original draft: CW, RR

Writing – review & editing: CW, EMJ, RR

## Competing interests

The authors declare that they have no competing interests.

## Data and materials availability

All data are available in the main text or the supplementary materials.

## Supplementary Materials

Materials and Methods

Figs. S1 to S11

Tables S1 to S2

References (38–45)

MDAR Reproducibility Checklist

## Materials and Methods

### *C. elegans* strains and maintenance

*C. elegans* were grown with standard procedures and maintained at 15°C on Nematode Growth Media (NGM) plates seeded with *E. coli* (OP50) (*38*). The strains used in this study can be found in table S1.

### Transgenes and transgenic animals

To make the DNA plasmid containing *rgef-1p::gfp::rab-7::rab-7 3’UTR*, the coding sequence and the 3’UTR of *rab-7* were amplified from genomic DNA using PCR and primers 5’-catgaagcttatgtcgggaaccagaaagaa-3’ and 3’-catgctgcaggcctcagcgactacatcctc-5’ and digested using HindIII and PstI and inserted into the pSK vector digested with HindIII and PstI. *rgef-1p* was amplified from plasmid pMR2093 containing *rgef-1p::aak-2::gfp* using PCR and primers 5’-tcgacggtatcgataagcttatccttttcattttgaactcaccag-3’ and 3’-aagcttatcgataccgtcgac-5’. *gfp* was amplified from plasmid pPD95.77 from the Fire Lab kit containing *gfp* using PCR and primers 5’-cggcatcgacgacgacgacgatgagtaaaggagaagaacttttca-3’ and 3’-tggcatggatgaactatacaaaatgtcgggaaccagaaagaa-5’. *rgef-1p* and *gfp* were inserted into a vector containing the sequence *rab-7* amplified using PCR and primers 5’-atgtcgggaaccagaaagaa-3’ and 3’-aagcttatcgataccgtcgac-5’ to create *rgef-1p::gfp::rab-7::rab-7 3’UTR* using Gibson assembly (NEB E2611). A full list of reagents and resources can be found in table S2.

To make the DNA plasmid containing *myo-3p::gfp::rab-7::rab-7 3’UTR*, *nhx-2p::gfp::rab-7::rab-7 3’UTR*, and *pgp-12p::gfp::rab-7::rab-7 3’UTR*, a 2.8 kb, 1.9 kb, and 3.3 kb DNA sequence upstream of each respective gene was amplified using PCR and cloned into the vector containing the sequence *gfp::rab-7::rab-7 3’UTR* amplified from plasmid *rgef-1p::gfp::rab-7::rab-7 3’UTR* using PCR and primers 5’-atgagtaaaggagaagaacttttca-3’ and 3’-aagcttatcgataccgtcgac-5’ were used to generate tissue-specific *gfp::rab-7::rab-7 3’UTR* by using Gibson assembly (NEB E2611).

For tissue-specific RNAi mutants, we used *rde-1(mkc36)* mutants where wild type expression of *rde-1* was rescued using tissue-specific promoters in the appropriate genetic backgrounds required for that specific experiment (*39*). Germline-specific and neuron-specific RNAi were validated in their original publications (*3, 39, 40*). Muscle-specific RNAi was generated for this study. The strains MR2283 and MR2337 were validated by feeding of dsRNA against *unc-112* (muscle), *egg-5* (germ line), or *unc-13* (neurons) and the phenotypes were scored. Each dsRNA treatment exhibits a unique phenotype, such as paralysis (*unc-112* RNAi), embryonic lethal (*egg-5* RNAi), or paralysis (*unc-13* RNAi). Muscle-specific RNAi only exhibited paralysis when fed dsRNA against unc-112.

To generate a DNA plasmid containing *pie-1p::his-15::gfp::3xmir-51::tbb-2 3’UTR*, the *tbb-2 3’UTR* sequence was amplified from genomic DNA using PCR and primers 5’-tggatgaactatacaaatagatgcaagatcctttcaagca-3’ and 3’-aggttttcaccgtcatcacccgcgaaaaacccatgtaagt-5’ and the vector containing the sequence of *pie-1p::his-15::gfp* amplified from plasmid containing *pie-1p::his-15::gfp::egg-6 3’UTR* using PCR and primers 5’-ggtgatgacggtgaaaacct-3’ and 3’-ctatttgtatagttcatccatgcc-5’ were used to generate *pie-1p::his-15::gfp::tbb-2 3’UTR* by using Gibson assembly (NEB E2611). To add the wild type or mutant *3xmir-51* seed sequences, *pie-1p::his-15::gfp::tbb-2 3’UTR* was amplified using PCR with primers containing overhangs for the *3xmir-51*sequences using primers 5’-atttttacgggtttttcttttcgattcatttttacgggtttatgcaagatcctttcaagc-3’ and 3’-gaatcgaaataggaaacccgtaaaaatgaatcgaaactatttgtatagttcatccatgcc-5’ for wild type, and primers 5’-attttaaggcgattttcttttcgattcattttaaggcgattatgcaagatcctttcaagc-3’ and 3’-gaatcgaaataggaatcgccttaaaatgaatcgaaactatttgtatagttcatccatgcc-5’ for mutant. The PCR product was subjected to T4 PNK (NEB M0201S) and T4 DNA ligase (NEB M0202S) to make *pie-1p::his-15::gfp::3xmir-51::tbb-2 3’UTR* with either wild type or mutant *3xmir-51* sites. The plasmids were cloned into the pCFJ151 MosSCI vector targeting the *ttTi5605* targeting region (Addgene) (*41*). The sequence of the scrambled seed sequence is as follows: 5’- TTTCGATTCATTTT<u>aAgGcGaTTccta</u>TTTCGATTCATTTT<u>aAgGcGaTTttct</u>TTTCGATTCATTTT<u>aAgGcGaT</u>T-3’. The seed sequences are underlined, and the sites of mutagenesis are in lowercase.

To generate a DNA plasmid containing *rgef-1p::mir-51::unc-54 3’UTR*, the pre-microRNA sequence of *mir-51* was amplified from genomic DNA using PCR and primers 5’-cggcatcgacgacgacgacggtccgaaaagtccgtctacc-3’ and 3’- cagttggaattctacgaatgaactgtattgctgctgggc-5’ and the vector containing the sequence of *rgef-1p* and *unc-54 3’UTR* amplified from plasmid *rgef-1p::aak-2::unc-54 3’UTR* using PCR and primers 5’-cattcgtagaattccaactgagc-3’ and 3’-cgtcgtcgtcgtcgatgc-5’ were used to generate *rgef-1p::mir-51::unc-54 3’UTR* by using Gibson assembly (NEB E2611).

To generate a DNA plasmid containing *rgef-1p::mir-51::unc-54 3’UTR* with scrambled EXOmotifs in the mature *mir-51* sequence, the *pre-mir-51* sequence was first inserted into a 3kb empty vector from pMR377 digested with XcmI. Then the three EXOmotifs on *pre-mir-51* were scrambled via site-directed mutagenesis using three sets of primers; set 1 primers 5’-ttactggtcaaaaagtgaacatgg-3’ and 3’- gaacgataggagctacgggtagac-5’, set 2 primers 5’-gtccgaagcaggtacaggtgca-3’ and 3’- ttcactttttgaccagtaagaacg-5’, and set 3 primers 5’-aagctggggctctggg-3’ and 3’- gaacaccctactcgccgtgc-5’. The resulting PCR product was treated with T4 PNK (NEB M0201S) and T4 DNA ligase (NEB M0202S), then verified through sequencing. Plasmids containing neuronally expressed wild-type or scrambled *mir-51* were injected at 1 ng μl^-1^. The sequence of the scrambled *mir-51* is as follows: 5’- GTCCGAAAAGTCCGTCTACCCGTAGCTCCTATCgttcTTACTGGTCAAAAAGTGAAgtccGAAGCAGGTACAGGTGCACGGCGAGTAGGGTgtccAAGCT-3’ with the sites of mutagenesis in lowercase.

To generate a DNA plasmid containing *rgef-1p::mKate::3xFLAG::alg-1*, the coding sequence of *alg-1* was amplified from genomic DNA using PCR and primers 5’- acaaggacgacgacgacaagatggaagaccaatggttgct-3’ and 3’- cagttggaattctacgaatgttaagcaaagtacatgacgttgttggc-5’ and the coding sequence of *mKate::3xFLAG* was amplified from a plasmid containing *mKate::3xFLAG* using PCR and primers 5’-cggcatcgacgacgacgacgatggtttccgagttgatcaagg-3’ and 3’- cttgtcgtcgtcgtccttgtagtcgatAtcgtggtccttgtagtcaccgtcgtggtccttgtagtccttacgatgtccgagcttgg-5’ and the vector containing *rgef-1p* and *unc-54 3’UTR* was amplified from plasmid *rgef-1p::aak-2::unc-54 3’UTR* using PCR and primers 5’-cattcgtagaattccaactgagc-3’ and 3’-cgtcgtcgtcgtcgatgc-5’ were used to generate *rgef-1p::mKate::3xFLAG::alg-1* by using Gibson assembly (NEB E2611). To generate a DNA plasmid containing *rgef-1p::mKate::3xFLAG::alg-2*, the same protocol was used but the coding sequence of *alg-1* was replaced with *alg-2* that was amplified from genomic DNA using PCR and primers 5’-acaaggacgacgacgacaagatgttccctctgcctgtacac-3’ and 3’-cagttggaattctacgaatgttaggcaaaatacatgacgttgttc-5’.

To generate a kinase dead variant of SID-3, aaa (lysine) was mutated into gca (alanine) in the MR1963 *daf-2; aak(0); tbc-7(rr166)* strain using CRISPR-Cas9 with sgRNA sequence 3’-cgacatttctccaaatattcaagacatctcgcaatag-5’ and a repair template/ultramer DNA oligonucleotide (Integrated DNA Technologies) containing the desired K139A mutation (*26*).

Transgenic animals were generated by injecting transgenes (15 ng μl^-1^ each) mixed with a co-injection marker pRF-4 (120 ng μl^-1^) (Addgene) expressing a dominant negative variant of the *rol-6* gene and an empty vector pSK (up to 200 ng μl^-1^ of DNA) (Addgene) using standard methods (*42*). To create transgenic animals with germline *his-15::gfp* expression, EG6699 (*41*) (*ttTi5605 II; unc-119(ed3) III; oxEx1578*) animals were injected with a target transgene (50 ng μl^-1^) with plasmid pJL43.1 (50 ng μl^-1^) (Addgene) that expresses MosTase under a germline promoter and plasmid pMR910 (20 ng μl^-1^) that expresses GFP under a pharyngeal promoter using standard methods (*42*).

### Synchronization

A population of genetically identical animals were synchronized using alkaline hypochlorite. The resulting embryos were allowed to hatch in the absence of food in M9 buffer before they were plated on NGM plates seeded with OP50 (alternatively, on HT115 for RNAi experiments). The plates were incubated in 25°C for 96 hours to induce dauer formation and to allow the animals to spend at least 48 hours in the dauer stage. Afterwards, the plates were switched back to 15°C to allow for recovery and resumption of normal development.

### Quantification of AMPK germline defects

A population of genetically identical animals were synchronized and plated on either NGM plates seeded with OP50 or HT115 expressing dsRNA according to the conditions specified in the figures then incubated at the restrictive temperature of 25°C to induce dauer formation for a total of 96 hours. The dauer larvae were then separate onto individual plates and switched into the permissive temperature of 15°C to trigger dauer exit. The post-dauer fertility was assessed after 7 days and animals were deemed fertile if they gave rise to viable progeny.

For quantification of the number of germ cells in the dauer larvae. Dauer larvae were washed off a plate after spending a total of 96 hours in the restrictive temperature of 25°C and soaked in Carnoy’s solution (60% ethanol, 30% acetic acid, 10% chloroform) overnight on a shaker. Afterwards, the Carnoy’s solution was removed, and the worm pellet was washed twice with 1x PBS with 0.1% Tween-20 (PBST). The larvae were then stained with 0.1 mg mL^-1^ DAPI (Roche 10236276001) for 30 minutes while shaking. Next, the DAPI solution was removed, and the worm pellet was washed four times with PBST. Germ cells per dauer gonad was determined based on their position and their nuclear morphology and quantified manually. The conditions and genotypes of the animals were not blinded.

### Confocal microscopy

All confocal images were taken and analysed as previously described (*2, 43*). Briefly, images were acquired in a stack of 56 z-planes in increments of 0.2 μm. The DAPI signal was acquired with a wide-field X-Cite 120 florescence illumination system (Excelitas Technologies), while the GFP and mKate signals were acquired with a Quorum WaveFX spinning disc confocal system (Quorum Technologies), both integrated with the Leica DMI6000B microscope with 63x oil-immersion objective. Maximum intensity projection was generated using NIH ImageJ. Each stack was spliced together with overlap to create images of whole animals. Worms were artificially straightened using NIH ImageJ. Scale bars were generated using NIH ImageJ. The conditions and genotypes of the animals were not blinded.

### Negative-stain and immunogold labelling transmission electron microscopy

Carbon-coated copper grids (Agar Scientific) were glow-discharged for 20 seconds at 20 mA before sample preparation using EasiGlow Glow Discharge Cleaning System (PELCO). 5 μl of isolated exosomes was applied to the grid and allowed to settle for 5 minutes. The grids were blocked with 5 μl of PBS containing 0.5% BSA for 10 min at room temperature. The grids were incubated with mouse anti-HA.11 epitope tag antibody (BioLegend, 901501, 1:1000, RRID: AB_2565006) for 30 min at RT, then washed with PBS 3 times for a total of 10 min. The grids were incubated with 6 nm Colloidal Gold AffiniPure Goat Anti-Mouse IgG (H+L) (Jackson ImmunoResearch, 115-195-146, 1:20, AB_2338728) for 30 min at RT, then washed with PBS 3 times for a total of 10 min. Grids were stained with 5 μl of 2% uranyl acetate (Electron Microscopy Sciences) for 45 seconds. The excess solution was blotted off the grid using filter paper after each step. Staining was finished at least 1 hour before imaging. The grids were imaged using a Talos^TM^ F200X G2 (S)TEM (Thermo Scientific).

### RNA interference

For RNAi experiments, a population of genetically identical animals were synchronized and allowed to hatch in M9 buffer. L1 larvae are then plated on NGM plates supplemented with 1 mM IPTG (BioShop IPT002) and 50 μg mL^-1^ ampicillin (Fisher BioReagents BP176025) seeded with HT115 either containing an empty vector L4440 or expressing dsRNA against a gene of interest. RNAi constructs were obtained from the Ahringer RNAi library (*44*). All RNAi clones were sequence verified. The conditions and genotypes of the animals and bacteria were not blinded.

### Soaking of *daf-2; aak(0)* dauer larvae with dsRNA

Gravid *daf-2* animals were synchronized and incubated at the restrictive temperature of 25°C to induce dauer formation. Animals were harvested after 48 hours at 25°C and subjected to TRIzol^TM^ Reagent (Invitrogen 15596026) for RNA isolation. The isolated RNA was incubated at 65°C for 10 min and allowed to reanneal at room temperature for another 10 min. The reannealed RNA was either used directly for dsRNA soaking or subjected to RNase A (Thermo Scientific EN0531) treatment according to manufacturer’s instruction for Fig. 2c. For Fig. 2b, the mRNA was separated from the isolated RNA using the Poly(A) mRNA Magnetic Isolation Module (NEB E7490S) and Magnetic Separation Rack (NEB S1507S) according to manufacturer’s protocol. Then, gravid *daf-2; aak(0)* animals were synchronized and the resulting embryos were soaked with dsRNA from *daf-2* dauer larvae, RNase inhibitor (Applied Biosystems, N8080119), and a drop of worm HB101 superfood, and placed in the restriction temperature of 25°C to induce dauer formation. The animals were kept in 25°C for 96 hours before being switched into the permissive temperature of 15°C to trigger dauer recovery. The post-dauer fertility was scored 7 days afterwards. The conditions and genotypes of the animals were not blinded.

### Western blot

For Fig. 4d, isolated exosomes were subjected to 1% Triton X-100 (Sigma-Aldrich X100), 50 μg mL^-1^ Proteinase K (Promega V3021), both, or neither then incubated at 60°C for 1 hr. For Fig. 4c, the isolated exosomes were not subjected to any treatment before Western blot analysis. The isolated exosomes were mixed with 5 μl of loading buffer (5% β-mercaptoethanol, 0.02% bromophenol blue, 30% glycerol, 10% sodium dodecyl sulfate, 250 mM pH 6.8 Tris-Cl) and heated at 100°C for 5 minutes. Protein concentration was determined using a NanoDrop™ 2000c Spectrophotometer (Thermo Scientific ND-2000).

For whole protein lysates from *C. elegans*, either 500 dauer larvae or 200 post-dauer adult animals were mixed with loading buffer and subjected to multiple rounds of freeze-boiling. Protein lysates were frozen in liquid nitrogen and boiled on a 100°C heat block for 4.5 minutes for at least 4 cycles prior to SDS-PAGE.

The exosome sample was separated using SDS-PAGE and transferred to a nitrocellulose membrane (Bio-Rad, 1620115) before subjected to immunoblotting. Western blot analysis was performed using anti-HA.11 epitope tag antibody (BioLegend, 901501, 1:1000, RRID: AB_2565006), anti-FLAG (Sigma-Aldrich, F7425, 1:1000, RRID: AB_439687), or anti-GFP (produced by Roy Lab, 1:1000). Membranes were incubated with horseradish-peroxidase-conjugated anti-rabbit (Bio-Rad, 1721019, 1:2000, RRID: AB_11125143) or anti-mouse (SouthernBiotech, 1036-05, 1:2000, RRID: AB_2794348) secondary antibodies and visualized using a MicroChemi (DNR Bio Imaging Systems) and GelCapture software (Version 7.0.18).

### Exosome isolation

*daf-2* animals expressing HA::GFP::RAB-7 and mKate::3xFLAG::ALG-1 in the neurons were synchronized and put in the restrictive temperature of 25°C to induce dauer formation and incubated for 96 hours. Dauer larvae were harvested by washing the plates with M9 buffer. The dauer larvae were lysed using a Dounce tissue grinder set (Kimble 885300-0007) in PBS with Halt™ Protease Inhibitor Cocktail (100X) (Thermo Scientific 78429) until no intact worms remained. The protein lysate was pre-cleared through centrifugation for 15 min at 13,500 rpm at 4°C. Exosomes were isolated using the exoEasy maxi kit (Qiagen 76064) according to manufacturer’s protocol. The integrity was the isolated exosomes was verified using negative stain TEM and Western blotting against RAB-7.

### Immunoprecipitation

Immunoprecipitation against HA-tagged exosomes was performed using Abcam Immunoprecipitation protocol on pan-exosomes isolated using the exoEasy kit (Qiagen). Protein G beads (Thermo Scientific) were used according to manufacturer’s instructions. Briefly, 10 mg of protein G agarose beads were crosslinked to 5 μL of mouse anti-HA antibody (BioLegend, 901501, 1:1000, RRID: AB_2565006) using an effective concentration of 6.5 mg mL^-1^ DMP. No antibody control beads did not contain any antibodies but was still subjected to the crosslinking process. The antibody-bead mixture was mixed with 400 μL of isolated exosomes overnight at 4°C and then subjected to three washes with PBS. 150 μL of beads were boiled with loading buffer (described above) for 10 mins and loaded into the SDS-PAGE. 40 μL of exosomes from the exosome isolation was loaded into the SDS-PAGE as an input control. A rabbit anti-GFP primary antibody was used to detect HA::GFP::RAB-7 and a rabbit anti-CD63/TSP-7 antibody (Abclonal, A5271, 1:500, RRID: AB_2766092) was used to detect CD63/TSP-7.

### Quantitative reverse transcriptase-polymerase chain reaction

Gravid animals were synchronized and incubated at the restrictive temperature of 25°C to induce dauer formation. Animals were harvested after 48 hours at 25°C and subjected to TRIzol^TM^ Reagent (Invitrogen 15596026) for RNA isolation. The concentration of the RNA was determined with a NanoDrop™ 2000c Spectrophotometer (Thermo Scientific ND-2000). The quality of the RNA was determined with a NanoDrop™ 2000c Spectrophotometer (Thermo Scientific ND-2000) and agarose gel electrophoresis. 1000 ng of purified RNA was used to synthesize cDNA using a High-Capacity RNA-to-cDNA^TM^ Kit (AppliedBiosystems, 4387406). The gene expression levels were determined by RT-qPCR using a 2x SyberGreen qPCR Master-mix (ZmTech Scientifique, Q2100N) and a Bio-Rad CFX384 Real-Time 96-well PCR qPCR Detection System (Bio-Rad). 10 ng of cDNA was used for each qPCR reaction. The gene expression data was analyzed using CFX Maestro Software (Bio-Rad). Relative gene expression was calculated by normalizing to the expression of an alpha-tubulin gene, *tba-1*, which as the loading control. The gene expression was repeated three times and the mean and SEM were plotted. The following primers were used for qPCR: *tba-1* (5’- TCACCAACAGTTGCTTCGAG-3’, 3’-ACGTCCTTTGGAACGACATC-5’), *spe-26* (5’- TTCTCATCATCATCGGTGGA-3’, 3’-GCTTCTGCTGGGACACTTTC-5’), *pro-2* (5’-TGCCACAATTAAAGCCATGA-3’, 3’-GACGACCGCCTTAATCAAAA-5’), *ppk-2* (5’- AACAATCGAGGGCTCAGAGA-3’, 3’-CCGCACGGGATACTGTAGAT-5’), *pmk-1* (5’- GGAACTGTTTGTGCTGCTGA-3’, 3’-TGTACGACGGGCATGAATTA-5’), *mek-2* (5’- ATTGATTCGATGGCCAACTC-3’, 3’-GTGGGATCCTGTGAGTCGTT-5’).

### Statistics

Bar graphs and fluorescence intensity graphs were generated through GraphPad Prism 7. Fluorescence intensity was measured using NIH ImageJ. Statistical tests were performed using GraphPad Prism 7 and XLSTAT plugin for Microsoft Excel. The types of statistical tests used in each experiment, n numbers, P values, and other related measures are indicated in each figure and the associated legend. In all figures, ****P < 0.0001, ***P < 0.001, **P < 0.01, *P < 0.05; ns, not significant. Replications were done by using different worms performed on different days. Replications for all imaging experiments were performed on different worms on different days. Representative images were generated from more than 10 worms in at least three independently generated lines. RNAi experiments were performed on independently cultured animals and bacterial cultures. Western blots were performed on independently isolated exosomes purifications or animals at least twice. For all *C. elegans* strain used in this study, a population of worms was grown together under identical conditions, and the worms were randomly distributed into different conditions. Worms from a population were randomly chosen for RNAi analyses, post-dauer fertility, Western blot analysis, and imaging experiments.

## Notes

### Competing Interest Statement

The authors have declared no competing interest.

