## Supplementary material for "Neuronal exosomes transport a miRNA/RISC cargo to preserve germline stem cell integrity during energy stress"

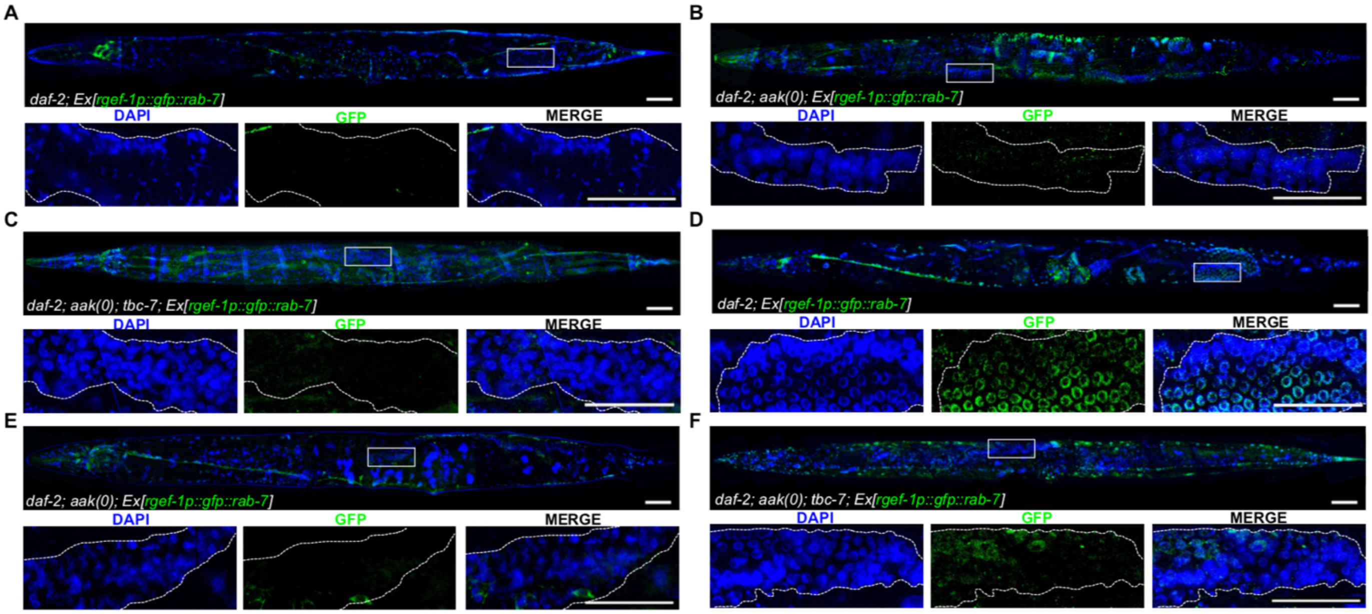

**Fig. S1.**

**Neuronal RAB-7 that localizes to the germ line during the dauer stage persists into the post-dauer germ line.** (**A-C**) Confocal images of neuronal GFP::RAB-7 expression in (A) *daf-2*, (B) *daf-2; aak(0)*, and (C) *daf-2; aak(0); tbc-7* adults that did not transit through the dauer stage. (**D-F**) Confocal images of neuronal GFP::RAB-7 expression in post-dauer adults in (D) *daf-2*, (E) *daf-2; aak(0)*, and (F) *daf-2; aak(0); tbc-7* mutants. Solid white frame indicates position of the higher magnification insets (below). Dotted white lines outline the gonad. Scale bars, 50 $\mu$m. Representative of three independently generated transgenic lines.

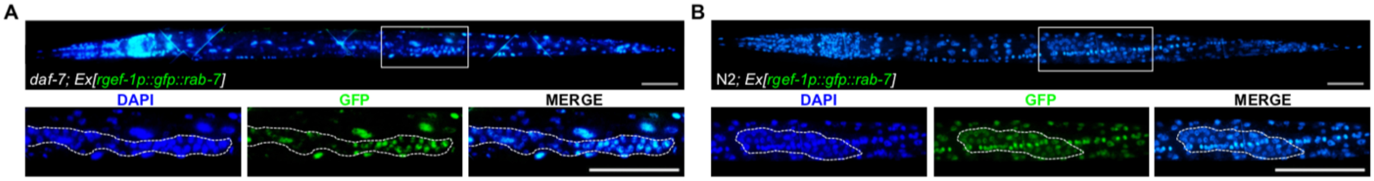

**Fig. S2.**

**The mechanism of neuron to germ line transfer of GFP::RAB-7 is conserved across different modes of dauer formation.** (**A-B**) Confocal images of neuronal GFP::RAB-7 expression in (A) *daf-7* and (B) N2 animals treated with dauer pheromone. Solid white frame indicates position of the higher magnification insets (below). Dotted white lines outline the gonad. Scale bars, 50 $\mu$m. Representative of three independently-generated transgenic lines.

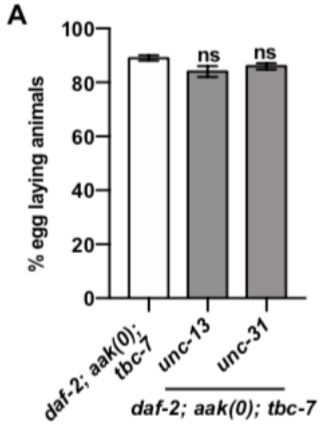

**Fig. S3.**

**Synaptic vesicle or dense-core vesicle secretion does not play a role in regulating germ cell integrity.** (A) Post-dauer fertility of *daf-2; aak(0); tbc-7* mutants that contain either a variant of *unc-13* (synaptic vesicle) with a premature stop codon or a deletion of *unc-31* (dense-core vesicle). Data is mean ± SEM. *P* values are derived from Marascuilo procedure (A); n = 50 animals per condition for post-dauer fertility. Post-dauer fertility data are representative of three independent experiments. ns, not significant.

**A**

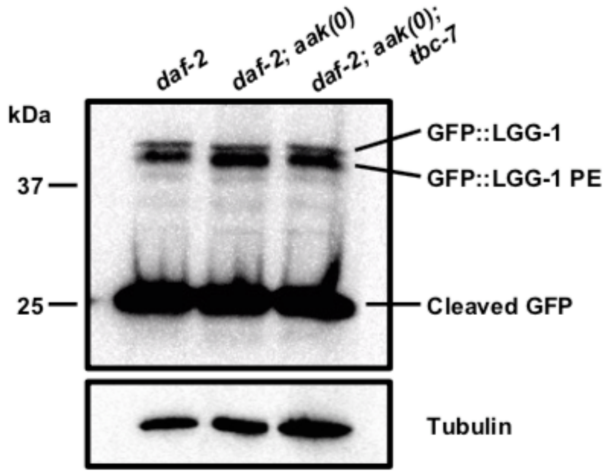

**Fig. S4.**

**Autophagic flux is not affected in *tbc-7*-suppressed mutants.** (**A**) Western analysis with anti-GFP antibodies on whole protein lysate of *daf-2*, *daf-2; aak(0)*, and *daf-2; aak(0); tbc-7* mutants expressing a GFP::LGG-1 transgene. The phosphatidylethanolamine (PE) conjugated LGG-1 variant and cleaved GFP are readouts of autophagic flux. Representative of two independent experiments.

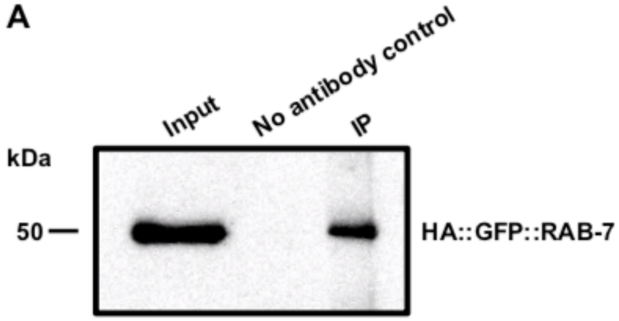

**Fig. S5.**
**Western analysis of immunoprecipitation of HA-tagged exosomes.** (**A**) An immunoprecipitation experiment using an antibody against HA was used to isolate HA::GFP::RAB-7 containing exosomes. Western analysis with an antibody against GFP was conducted to verify the efficacy and specificity of the immunoprecipitation. The input is purified pan-exosomes from *daf-2* mutants expressing neuronal HA::GFP::RAB-7. The no antibody control does not contain any anti-HA antibody but was subjected to the same crosslinking procedure with the beads. The IP was performed using protein G beads crosslinked with anti-HA antibodies on purified pan-exosomes and eluted through boiling with loading buffer.

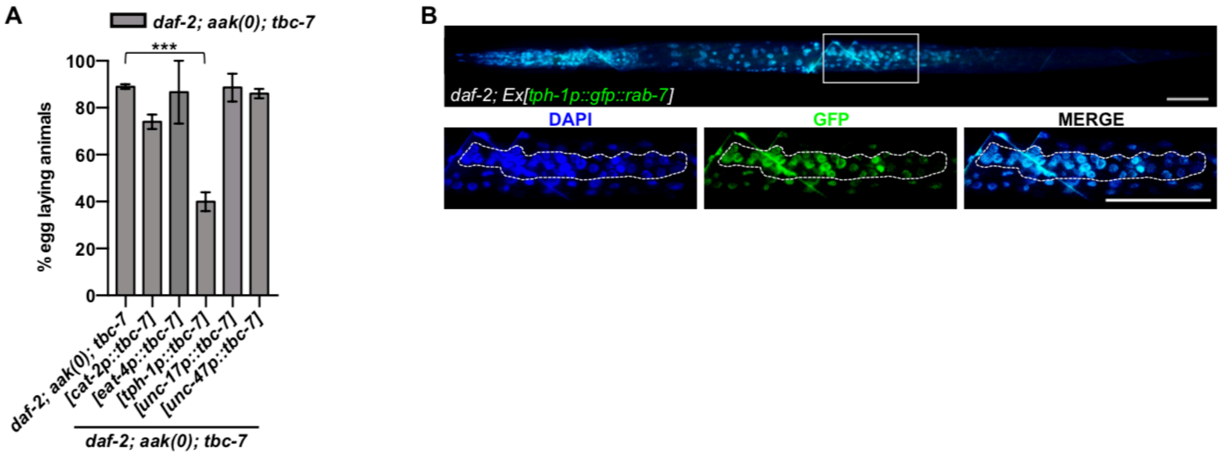

**Fig. S6.**

**Serotonergic neurons are responsible for the production and secretion of GFP::RAB-7 exosomes during the dauer stage.** (**A**) Post-dauer fertility of *daf-2; aak(0); tbc-7* mutants with wild type *tbc-7* expressed in dopaminergic (*cat-2p*), glutaminergic (*eat-4p*), serotonergic (*tph-1p*), cholinergic (*unc-17p*), and GABAergic (*unc-47p*) neurons. Data is mean ± SEM. *P* values are derived from Marascuilo procedure (A). ***P < 0.0001. Post-dauer fertility data are representative of three independent experiments. n = 50 animals per condition for post-dauer fertility. (**B**) Confocal images of GFP::RAB-7 expression in dauer larvae expressed in serotonergic neurons (*tph-1p*). Solid white frame indicates position of the higher magnification insets (below). Dotted white lines outline the gonad. Scale bars, 50 $\mu$m. Representative of three independently-generated transgenic lines.

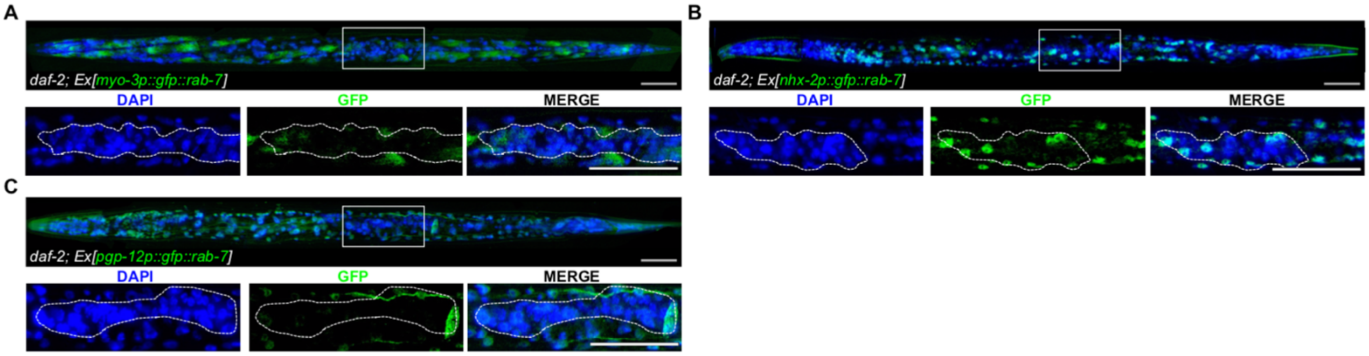

**Fig. S7.**

**The localization of the RAB-7-associated exosomes to the germ line requires that they be formed in the neurons.** (**A-C**) Confocal images of GFP::RAB-7 expression in dauer larvae expressed in the (A) muscle (*myo-3p*), (B) intestine (*nhx-2p*), or (C) the excretory system (*pgp-12p*). Solid white frame indicates position of the higher magnification insets (below). Dotted white lines outline the gonad. Scale bars, 50 $\mu$m. Representative of three independently generated transgenic lines.

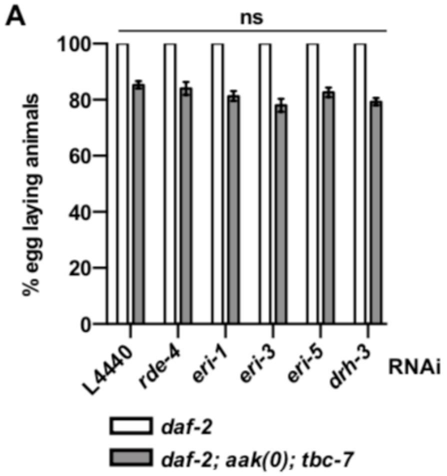

**Fig. S8.**
**Other classes of small RNAs are not required to maintain germ cell integrity in post-dauer AMPK mutants.** (**A**) Post-dauer fertilities of *daf-2* and *daf-2; aak(0); tbc-7* mutants subjected to RNAi of gene products involved in the biogenesis of ERGO-1 class 26G RNA and ALG-3/4 class 26G RNA. Data is mean ± SEM. *P* values are derived from Marascuilo procedure (A). ns, not significant. Post-dauer fertility data are representative of three independent experiments. n = 50 animals per condition for post-dauer fertility.

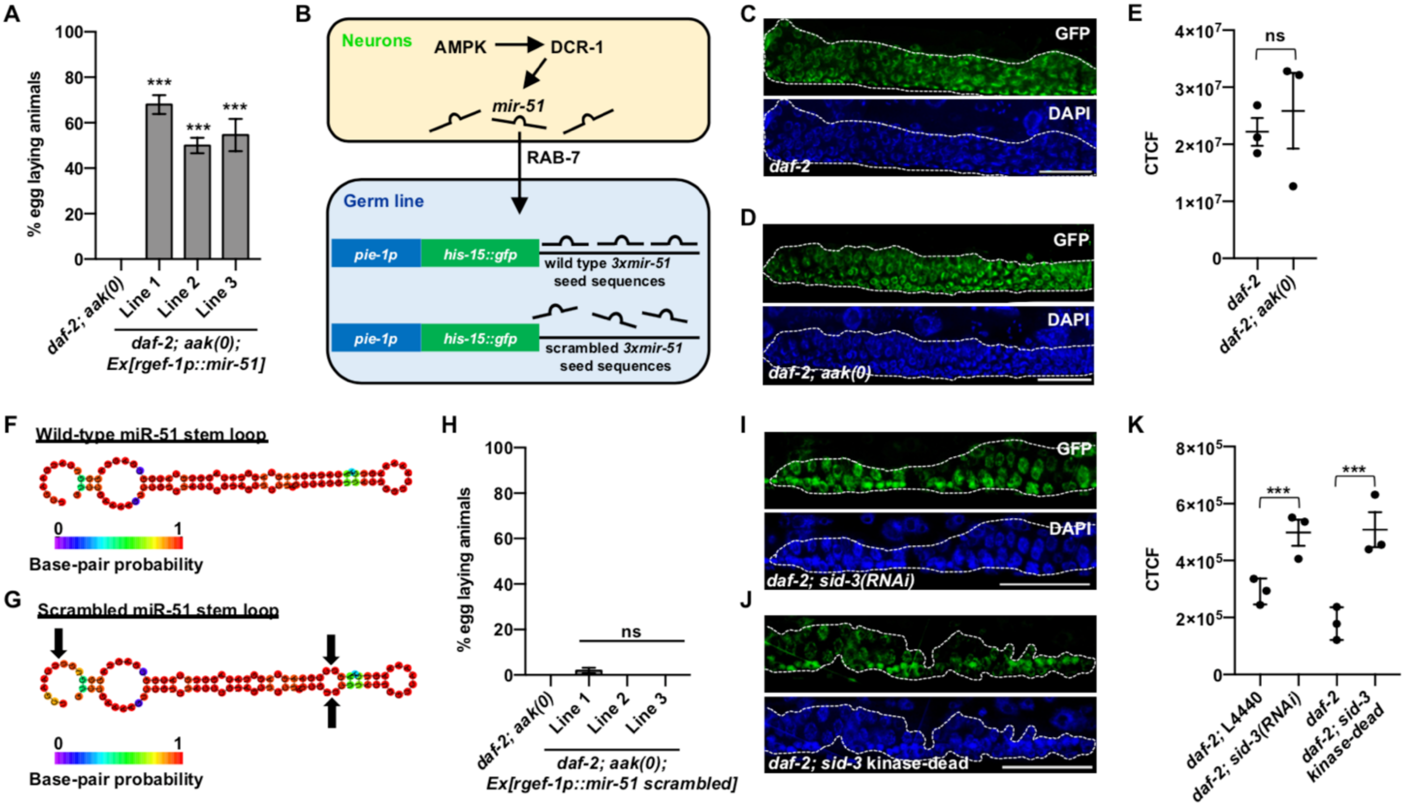

**Fig. S9.**

**Neuronally-expressed miRNAs are loaded into exosomes through EXOmotifs to regulate germ cell integrity non-autonomously.** (**A**) Post-dauer fertility of *daf-2; aak(0)* mutants expressing neuronal *mir-51*. (**B**) Graphic description of the *mir-51*-specifc germline miRNA sensor. (**C-D**) Confocal images of the miRNA sensor in (C) *daf-2* and (D) *daf-2; aak(0)* adult animals grown in replete conditions. (**E**) Corrected total cell fluorescence (CTCF) of GFP intensity by genotype. (**F-G**) Predicted structure for the hairpin miRNA for (F) wild-type or (G) EXOmotif scrambled *mir-51* by RNAfold Webserver software. Red or blue colouring indicate a high or low probability of pairing, respectively. Black arrows indicate the location of the mutated EXOmotif. (**H**) Post-dauer fertility of *daf-2; aak(0)* mutants expressing neuronal *mir-51* with scrambled EXOmotifs. (**I-J**) Confocal images of the miRNA sensor containing three wild-type *mir-51* seed sequences in *daf-2* dauer larvae (I) fed with bacteria expressing dsRNA that corresponds to *sid-3* or (J) with a CRISPR-engineered kinase-dead SID-3 K139A variant. (**K**) Corrected total cell fluorescence (CTCF) of GFP intensity by genotype. Solid white frame indicates position of the higher magnification insets (below). Dotted white lines outline the gonad. Scale bars, 50 $\mu$m. Representative of three independently-generated transgenic lines. *P* values are derived from Marascuilo procedure (A, H) and one-way ANOVA (E, K). Data is mean ± SEM. Post-dauer fertility data are representative of three independent experiments; n = 50 animals per condition. CTCF data is representative of three independent experiments; n = 15 animals per condition. ***$P < 0.0001$. ns, not significant.

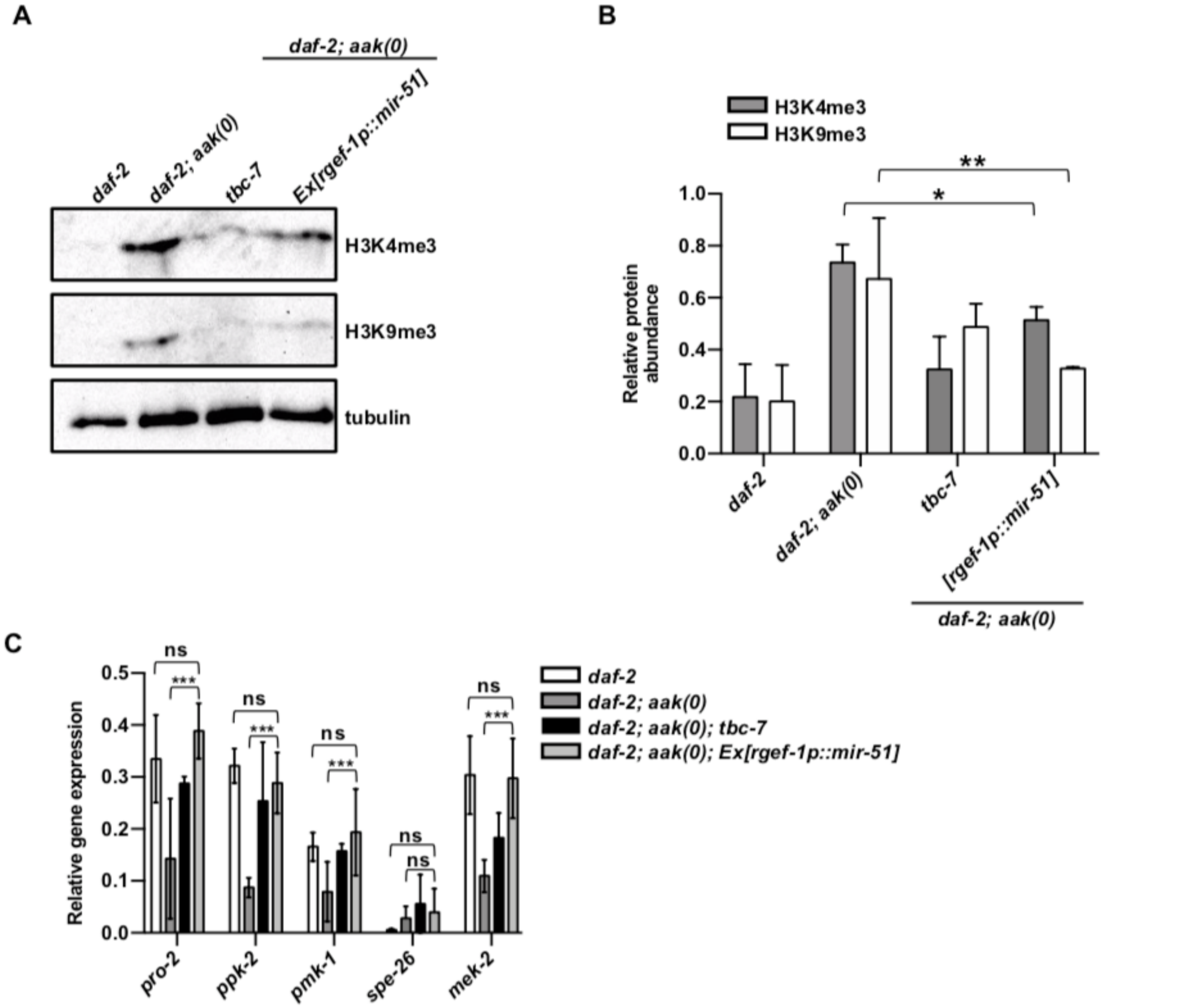

**Fig. S10.**

**Neuronal expression of *mir-51* corrects the aberrant chromatin marks and maladaptive germline gene expression characteristic of AMPK mutants.** (**A**) Global levels of H3K4me3 and H3K9me3 were quantified by performing whole-animal western analysis on dauer larvae. (**B**) Levels of chromatin marks were quantified and normalized to tubulin using ImageJ software. **P < 0.001, *P < 0.01 using Student's t-test. Representative of three independent experiments. (**C**) RT-qPCR was conducted on *daf-2; aak(0); Ex[rgef-1::mir-51]* mutants to quantify the levels of germline genes that were previously shown to be differentially expressed in the dauer stage. The expression between genotypes were normalized to *tba-1*. ***P < 0.001, ns, not significant using two-way ANOVA. Data is mean ± SEM. RT-qPCR data are representative of three independent experiments.

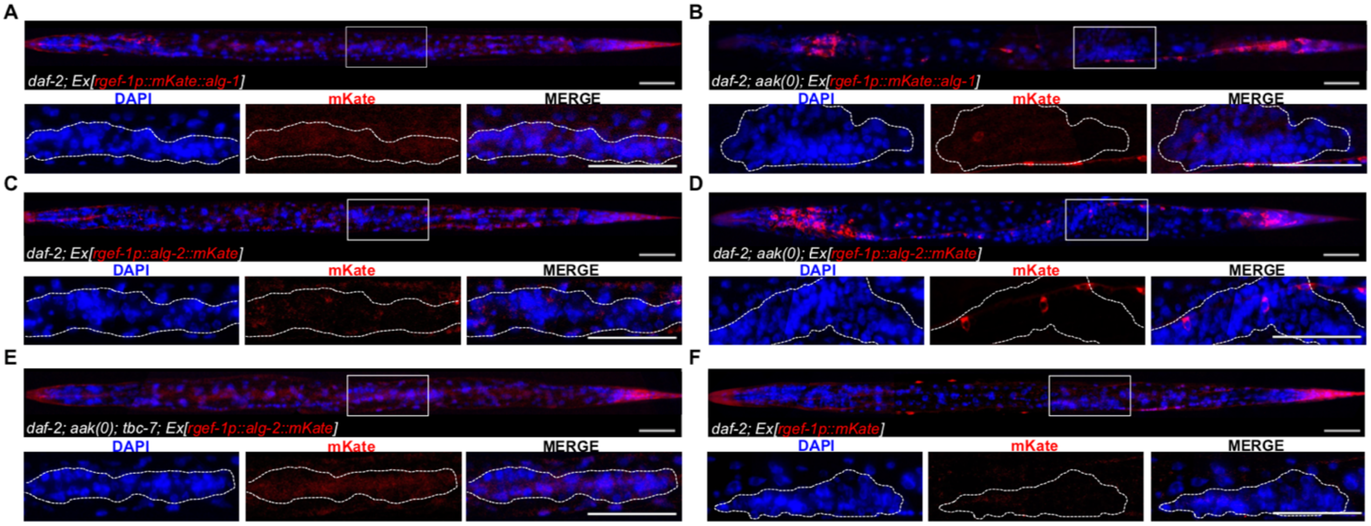

**Fig. S11.**

**Neuronally-expressed miRNA Argonautes are incorporated into exosomes in a RAB-7-dependent manner.** (**A-B**) Confocal images of neuronal mKate::ALG-1 expression in dauer larvae in (A) *daf-2* and (B) *daf-2; aak(0)*. (**C-E**) Confocal images of neuronal ALG-2::mKate expression in dauer larvae in (C) *daf-2*, (D) *daf-2; aak(0)*, and (E) *daf-2; aak(0); tbc-7* mutants. (**F**) Confocal image of neuronal mKate expression in *daf-2* dauer larvae. Solid white frame indicates position of the higher magnification insets (below). Dotted white lines outline the gonad. Scale bars, 50 $\mu$m. Representative of three independently-generated transgenic lines.

**Table S1.** ***Caenorhabditis elegans*** **strains from this study.**

| **Experimental Models: *Caenorhabditis elegans* Strains** | | |
|---|---|---|
| CB1370 | *daf-2(e1370)III* | (*45*) |
| MR1000 | *daf-2(e1370) aak-1(tm1944)III; aak-2(ok523)X* | (*2*) |
| MR1963 | *daf-2(e1370) aak-1(tm1944)III; tbc-7(rr166) aak-2(ok523)X* | (*3*) |
| MR2733 | *daf-2(e1370)III; rrEx761[rgef-1p::HA::gfp::rab-7; sur-5p::NeoR; rgef-1p::ollas::tsp-7; rgef-1p::3xFLAG::mKate::alg-1; rol-6]* | This study |
| MR2456 | *daf-2(e1370) aak-1(tm1944)III; aak-2(ok524)X; rrEx538[rgef-1p::gfp::rab-7]* | This study |
| MR2406 | *daf-2(e1370) aak-1(tm1944)III; tbc-7 (rr166) aak-2(ok524)X; rrEx535[rgef-1p::gfp::rab-7*] | This study |
| MR2289 | *mir-1(gk276)I; daf-2(e1370)III* | This study |
| MR2592 | *nDf49[mir-44; ZK930.12; mir-42; ZK930.15; mir-43]II; daf-2(e1370)III* | This study |
| MR2593 | *mir-1(gk276)I; nDf49[mir-44; ZK930.12; mir-42; ZK930.15; mir-43]II; daf-2(e1370)III* | This study |
| MR2283 | *mkcSi13[sun-1p::rde-1::sun-1 3'UTR + unc-119(+)]II; daf-2(e1370) aak-1(tm1944)III; rde-1(mkc36)V; tbc-7(rr166) aak-2(ok524)X* | This study |
| MR2337 | *daf-2(e1370) aak-1(tm1944)III; rde-1(mkc36)V; tbc-7(rr166) aak-2(ok523)X; rrEx527[rgef-1p::sid-1; rgef-1p::rde-1; rol-6]* | This study |
| MR2861 | *rrIs62[pie-1p::his-15::gfp::3xmir-51::tbb-2 3'UTR]II; unc-119(+)II; daf-2(e1370) unc-119(ed3)III; oxEx1678* | This study |
| MR2866 | *rrIs62[pie-1p::his-15::gfp::3xmir-51::tbb-2 3'UTR]II; unc-119(+)II; daf-2(e1370) aak-1(tm1944) unc-119(ed3)III; aak-2(ok524)X; oxEx1678* | This study |
| MR2462 | *daf-2(e1370) aak-1(tm1944)III; tbc-7(rr166) sid-3(rr301) aak-2(ok524)X* | This study |
| MR2298 | *daf-2(e1370) aak-1(tm1944)III; unc-31(e928)IV; tbc-7(rr166) aak-2(ok524)X* | This study |
| MR2299 | *unc-13(e1091)I; daf-2(e1370) aak-1(tm1944)III; tbc-7(rr166) aak-2(ok524)X* | This study |
| MR2726 | *daf-2(e1370)III; rrEx754[myo-3p::gfp::rab-7; rol-6]*), MR2724 (*daf-2(e1370)III; rrEx752[pgp-12p::gfp::rab-7; rol-6]*) | This study |
| MR2867 | *daf-2(e1370)III; rrEx875[nhx-2p::gfp::rab-7; rol-6]* | This study |
| MR2868 | *daf-2(e1370) aak-1(tm1944)III; aak-2(ok523)X; rrEx876[rgef-1p::mir-51; rol-6]* | This study |
| MR2862 | *rrIs62[pie-1p::his-15::gfp::3xmir-51::tbb-2 3'UTR]II; unc-119(+)II; daf-2(e1370) unc-119(ed3)III; mir-51(n4473)IV; oxEx1578* | This study |
| MR2869 | *rrIs62[pie-1p::his-15::gfp::3xmir-51::tbb-2 3'UTR]II; daf-2(e1370)III; mir-51(n4473)IV; rrEx877[rgef-1p::mir-51 wild-type]* | This study |

| | | |
|---|---|---|
| MR2870 | *rrIs62[pie-1p::his-15::gfp::3xmir-51::tbb-2 3'UTR]II; daf-2(e1370)III; mir-51(n4473)IV; rrEx878[rgef-1p::mir-51 scrambled; rol-6]* | This study |
| MR2863 | *rrIs63[pie-1p::his-15::gfp::scrambled 3xmir-51::tbb-2 3'UTR]II; unc-119(+)II; daf-2(e1370) unc-119(ed3)III; oxEx1678* | This study |
| MR2858 | *daf-2(e1370) aak-1(tm1944)III; aak-2(ok523)X; rrEx870[rgef-1p::mir-51 scrambled; rol-6]* | This study |
| MR2740 | *daf-2(e1370)III; rrEx767[rgef-1p::HA::gfp::rab-7; sur-5p::NeoR; rgef-1p::ollas::tsp-7; rgef-1p::alg-2::3xFLAG::mKate; rol-6]* | This study |
| MR2798 | *daf-2(e1370) aak-1(tm1944)III; aak-2(ok523)X; rrEx838[rgef-1p::HA::gfp::rab-7; sur-5p::NeoR; rgef-1p::ollas::tsp-7; rgef-1p::alg-2::3xFLAG::mKate; rol-6]* | This study |
| MR2815 | *daf-2(e1370) aak-1(tm1944)III; tbc-7(rr166) aak-2(ok523)X; rrEx847[rgef-1p::HA::gfp::rab-7; sur-5p::NeoR; rgef-1p::ollas::tsp-7; rgef-1p::alg-2::3xFLAG::mKate; rol-6]* | This study |
| MR2695 | *daf-2(e1370) III; rrEx847[rgef-1p::gfp::rab-7; rgef-1p::mKate; rol-6]* | This study |
| MR2871 | *daf-2(e1370) aak-1(tm1944)III; aak-2(ok523)X; rrEx879[rgef-1p::HA::gfp::rab-7; sur-5p::NeoR; rgef-1p::ollas::tsp-7; rgef-1p::3xFLAG::mKate::alg-1; rol-6]* | This study |
| MR2820 | *daf-2(e1370) aak-1(tm1944)III; tbc-7(rr166) aak-2(ok523)X; rrEx849[rgef-1p::HA::gfp::rab-7; sur-5p::NeoR; rgef-1p::ollas::tsp-7; rgef-1p::3xFLAG::mKate::alg-1; rol-6]* | This study |
| MR2287 | *daf-2(e1370)III; adIs2122[lgg-1::GFP rol6(df)]* | This study |
| MR2285 | *daf-2(e1370) aak-1(tm1944)III; aak-2(ok524)X; adIs2122[lgg-1::GFP rol6(df)]* | This study |
| MR2286 | *daf-2(e1370) aak-1(tm1944)III; tbc-7(rr166) aak-2(ok524)X; adIs2122[lgg-1::GFP rol6(df)]* | This study |
| MR2939 | *daf-7(e1372)III; rrEx917[rgef-1::HA::gfp::rab-7; sur-5::NeoR; rgef-1::ollas::tsp-7; rgef-1::3xFLAG::mKate; rol-6] line 1* | This study |
| MR2944 | *N2; rrEx[rgef-1p::HA::gfp::rab-7; rol-6]* | This study |
| MR2945 | *daf-2(e1370)III; rrEx923[tph-1p::HA::gfp::rab-7; rol-6]* | This study |
| MR2942 | *daf-2(e1370) aak-1(tm1944)III; tbc-7(rr166) aak-2(ok524)X; rrEX920[tph-1p::tbc-7; rol-6] line 1* | This study |
| MR2894 | *daf-2(e1370) aak-1(tm1944)III; tbc-7(rr166) aak-2(ok524)X; rrEx894[unc-17p::tbc-7] line 1* | This study |
| MR2896 | *daf-2(e1370) aak-1(tm1944)III; tbc-7(rr166) aak-2(ok524)X; rrEx896[eat-4p::tbc-7] line 1* | This study |
| MR2900 | *daf-2(e1370) aak-1(tm1944)III; tbc-7(rr166) aak-2(ok524)X; rrEx899[cat2p::tbc-7; rol-6] line 1* | This study |

| | | |
|---|---|---|
| MR2902 | *daf-2(e1370) aak-1(tm1944)III; tbc-7(rr166) aak-2(ok524)X; rrEx901[unc-48p::tbc-7; rol-6] line 1* | This study |

**Table S2. Reagents and resources from this study.**

| REAGENT or RESOURCE | SOURCE | IDENTIFIER |
|---|---|---|
| **Antibodies** | | |
| Mouse anti-HA.11 epitope tag antibody<br>Use 1:1000 for immunoblotting<br>Use 1:50 for transmission electron microscopy | BioLegend | 901501 |
| 6 nm Colloidal Gold AffiniPure Goat Anti-Mouse IgG (H+L)<br>Use 1:20 for transmission electron microscopy | Jackson ImmunoResearch | 115-195-146 |
| Rabbit anti-FLAG antibody<br>Use 1:1000 for immunoblotting | Sigma-Aldrich | F7425 |
| Rabbit anti-GFP antibody<br>Use 1:1000 for immunoblotting | This study | N/A |
| Horseradish-peroxidase-conjugated anti-rabbit secondary antibody<br>Use 1:2000 for immunoblotting | Bio-Rad | 1721019 |
| Horseradish-peroxidase-conjugated anti-mouse secondary antibody<br>Use 1:2000 for immunoblotting | SouthernBiotech | 1036-05 |
| Rabbit anti-CD63/TSP-7 antibody<br>Use 1:500 for immunoblotting | ABclonal | A52471 |
| **Bacterial Strains** | | |
| *Escherichia coli*: OP50 | CGC | N/A |
| *Escherichia coli*: HT115 | CGC | N/A |
| *Escherichia coli*: HB101 | CGC | N/A |
| **Chemicals, Enzymes, and Recombinant Proteins** | | |
| T4 PNK | NEB | M0201S |
| T4 DNA ligase | NEB | M0202S |
| DAPI | Roche | 10236276001 |
| Uranyl acetate | Electron microscopy sciences | NC1375332 |
| Isopropyl β- d-1-thiogalactopyranoside (IPTG) | BioShop | IPT002 |
| Ampicillin | Fisher BioReagents | BP176025 |
| TRIzol$^{TM}$ Reagent | Invitrogen | 15596026 |
| RNase A | Thermo Scientific | EN0531 |
| Poly(A) mRNA Magnetic Isolation Module | NEB | E7490S |

| | | |
|---|---|---|
| RNase inhibitor | Applied Biosystems | N8080119 |
| Triton X-100 | Sigma-Aldrich | X100 |
| Proteinase K | Promega | V3021 |
| Halt™ Protease Inhibitor Cocktail (100X) | Thermo Scientific | 78429 |
| Protein G-agarose beads | Millipore | P7700 |
| **Critical Commercial Assays** | | |
| Gibson Assembly® Master Mix | NEB | E2611 |
| exoEasy maxi kit | Qiagen | 76064 |
| High-Capacity RNA-to-cDNA™ Kit | AppliedBiosystems | 4387406 |
| 2x SyberGreen qPCR Master-mix | ZmTech Scientifique | Q2100N |
